# Structural attributes and principles of the neocortical connectome in the marmoset monkey

**DOI:** 10.1101/2020.02.28.969824

**Authors:** Panagiota Theodoni, Piotr Majka, David H. Reser, Daniel K. Wójcik, Marcello G.P. Rosa, Xiao-Jing Wang

## Abstract

The marmoset monkey has become an important primate model in Neuroscience. Here we characterize salient statistical properties of inter-areal connections of the marmoset cerebral cortex, using data from retrograde tracer injections. We found that the connectivity weights are highly heterogeneous, spanning five orders of magnitude, and are log-normally distributed. The cortico-cortical network is dense, heterogeneous and has high specificity. The reciprocal connections are the most prominent and the probability of connection between two areas decays with their functional dissimilarity. The laminar dependence of connections defines a hierarchical network correlated with microstructural properties of each area. The marmoset connectome reveals parallel streams associated with different sensory systems. Finally, the connectome is spatially embedded with a characteristic length that obeys a power law as a function of brain volume across species. These findings provide a connectomic basis for investigations of multiple interacting areas in a complex large-scale cortical system underlying cognitive processes.

## Introduction

Cognitive processes involve multiple interacting brain areas; however, characterization of the underlying inter-areal architecture is not yet fully understood. The last decade has seen a rapid change in neuroanatomy, from descriptive studies focused on few areas and nuclei at a time to those aimed at identifying the organizational principles, based on comprehensive and quantified large-scale datasets. Endeavors towards characterizing the full matrix of connections in the mouse brain using cellular resolution tracers are well underway^1–3^. In parallel, analogous attempts based on magnetic resonance imaging have enabled studies of cortical parcellation and structure-function relationships in the human brain connectivity, albeit at lower resolution^4^.

Many of the brain areas in humans that are involved in high-order cognitive processes and are affected in psychiatric conditions have no obvious rodent counterpart^5^. Primates have a large portion of the cortex devoted to vision, including many areas devoted to fine recognition of objects and to the complex spatial analyses required for oculomotor coordination^6^. The auditory cortex is similarly specialized, including a network of areas for identifying and localizing vocalizations^7^, while the motor cortex contains a unique mosaic of premotor areas for planning and executing movements^8^. Furthermore, the prefrontal cortex has expanded and become more complex through biological evolution. Therefore, to facilitate translation of discoveries in animal models to improvements in human health, studies of non-human primates are crucial to fill the gap between rodent and human models.

Macaques are the genus for which the most comprehensive knowledge of the connectional network of the cortex has been achieved, initially by studies based on meta-analyses of the literature, and more recently by retrograde tracer injections obtained with a consistent methodology^9^. Analyses of macaque data have already highlighted putative organizational principles of the primate cortical mesoscale connectome, enabling computational models to exhibit functional properties^10–12^. Similar data acquisition studies have also been achieved in the mouse brain^13,14^. However, extrapolating from any single species to human is problematic without knowledge of the scaling rules that govern anatomical similarities and differences^15^.

The marmoset is a non-human primate model with characteristics that complement those of the macaque in terms of facilitating analyses of brain anatomy, development, and function. Marmosets have a relatively short maturation cycle, which facilitates the development of transgenic lines and studies across the life span^16^. At the same time, the key anatomical features that motivate studies of the macaque brain are present, including well developed networks of frontal, posterior parietal and temporal association cortex. The volume of the marmoset brain is approximately 12 times smaller than that of the macaque brain, which in turn is 15 times smaller than the human brain, offering potential insights on scaling properties of the cortical network.

Here we provide the first account of the statistical properties of the marmoset cortical connectome, taking advantage of an online database of the results of retrograde tracer injections into 55, out of the 116 in total, cortical areas^17^. The dataset consists of connectivity weights, laminar origin of the projections and wiring distances. This allowed us to explore the statistical properties of the cortico-cortical connections, the architecture of the connectome by defining its hierarchical organization, and the characteristics of its spatial embedding. Furthermore, we studied how microstructural properties within each cortical area relate to the hierarchical organization of the connectome, providing a direct link to different scales within the cortex. In addition, we note conserved properties of the cortico-cortical connections across species, as well as differences that are species, or brain size, dependent. Finally, we present an allometric scaling law of the spatial localization of the connections with brain size which enables us to extrapolate this connectional attribute to humans.

## Results

### Connectivity weights are highly heterogeneous and log-normally distributed

We have analyzed the results of 143 retrograde tracer injections placed in 52 young adult marmosets (1.3 - 4.7 years; 31 male, 21 female; Supplementary Tables 1,2; Online Methods), available through the Marmoset Brain Connectivity Atlas (http://marmosetbrain.org)^17^. The injections were centered in 55 cortical areas (here referred to as *target areas*), which were distributed across the marmoset cortex (Fig. 1a; Supplementary Table 3). The use of retrograde tracers allows quantification of the number of neurons that project from 115 potential source areas to a given target area.

**Fig. 1.**
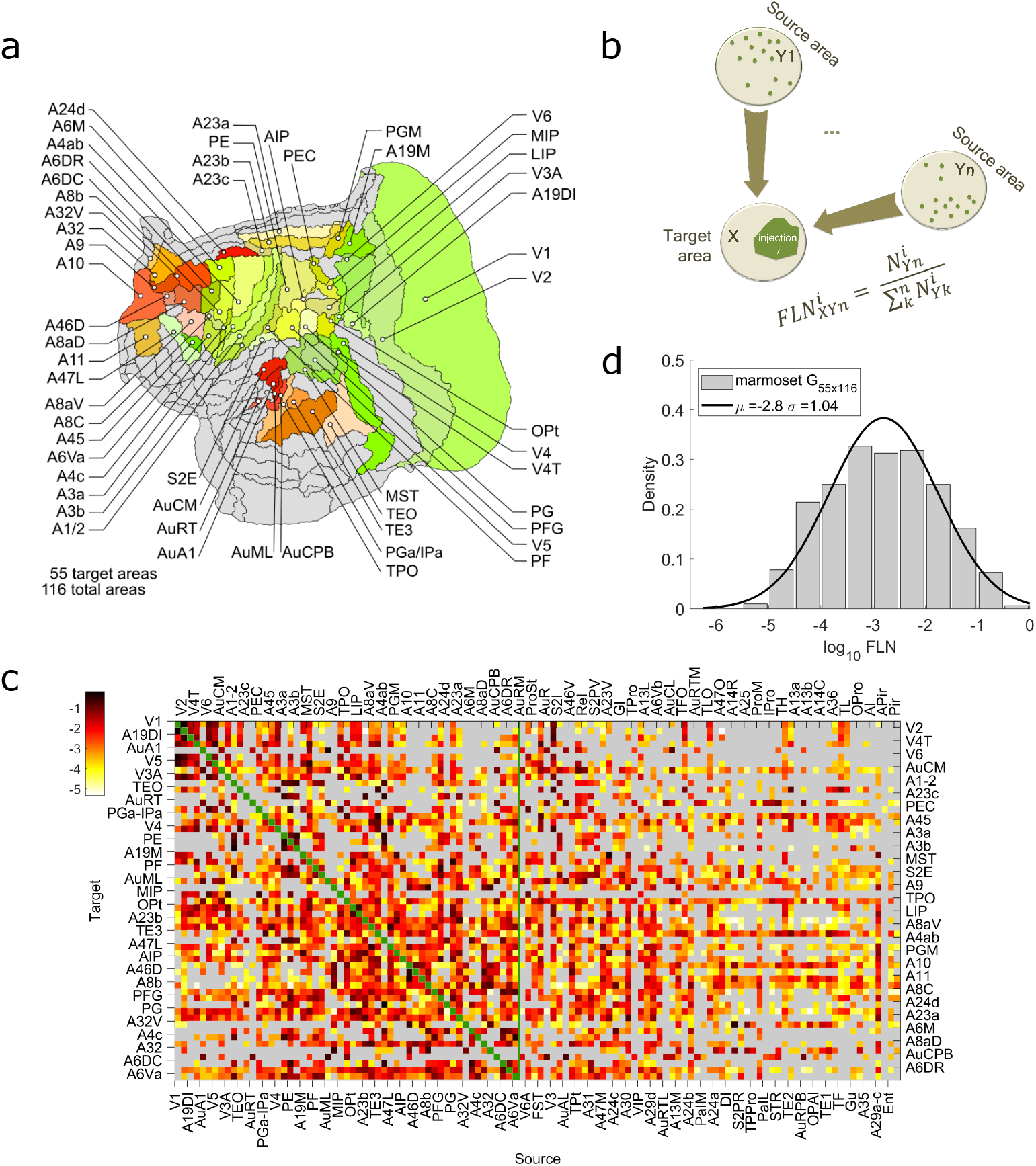
Cortico-cortical connectivity weights. **(a)** The analysis is focused on 55 cortical areas highlighted in different colors on the two-dimensional flattened map of the marmoset cortex. The grey shaded areas are those for which no tracer injection was available. **(b)** Schematic description of the fraction of labeled neurons found in area *Yn* after the retrograde tracer injection i in the cortical area 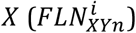. **(c)** The weighted and directed marmoset cortical interareal connectivity matrix. The rows are the 55 target areas and the columns the 116 source areas that provide inputs to the target areas. Each entry in the matrix is the base 10 logarithm of the arithmetic mean of the fraction of labeled neurons (*log*_10_*FLN*) across injections within the same target area. Grey: absence of connections, green, along the diagonal line: presence of intra-area connections (they have not been quantitively measured and the corresponding *FLN* is set to 0). The vertical green line defines the limit of the edge complete 55×55 subnetwork in which all inputs and outputs are known. **(d)** The distribution of the connectivity weights, shown in (c), reveals that the connectivity weights are highly heterogenous, they span five orders of magnitude, and they are log-normally distributed. Bin size = 0.5 on logarithmic scale. The black line is Gaussian fit to the *log*_10_*FLN* values.

A quantitative measure of the connectivity weight from each source area to a target area (Fig. 1b) is defined as the number of projection neurons found in each source area divided by the total number found across all source areas in the same hemisphere, called the fraction of labeled neurons (FLN). This analysis, which excluded connections from cells located in the same cytoarchitectural area (intrinsic connections), resulted in a 55×116 connectivity matrix (Fig. 1c). We found that the marmoset connectivity weights are highly heterogeneous, spanning five orders of magnitude, and are log-normally distributed (Fig. 1d), similar to macaque monkey^9^.

### The connectome is dense, heterogeneous, and has high specificity

In the graph theory framework, the cortex can be considered as a network where nodes correspond to areas, and edges to the connections between them. To characterize the network properties of the marmoset cortex we considered the edge-complete (*N*×*N* = 55)subnetwork, for which all inputs and outputs are known. This corresponds to approximately half of the full mesoscale connectome of this species (55/116 = 47.41%).

The inter-areal network density *ρ* = *M*/*N*(*N* − 1) defined as the fraction of existing connections (*M*)to all possible ones was found to be 62.43%. Even though the network is dense (Fig. 2a), there is high heterogeneity in the number of inputs and outputs of an area, as shown by its broad in- and out-degree normal distributions (Fig. 2b). Early analyses of cortico-cortical connectivity emphasized reciprocity of connections as a prominent property^18^. In the edge complete network of the marmoset connectome 50.3% are reciprocal connections, 24.24% are unidirectional and 25.45% are absent in both directions (similarly in macaque with densities 52.71%, 26.26%, and 24.26% correspondingly). While reciprocal connections are the most abundant, as well as stronger on average than the unidirectional (Supplementary Fig. 1a), bidirectionally absent connections are overrepresented when compared to those in an average random network that has same in- and out-degrees (and hence density), while unidirectional connections are underrepresented (Fig. 2c, left). Similar conclusions are reached when 3-node motifs, which are basic network building blocks^19^, are considered (Supplementary Fig. 1b). The motifs that are overrepresented in the marmoset connectome are those that include reciprocally present and absent connections (Fig. 2c, right). In addition, these appear more often (data/random ratio >2) than 2-node motifs (data/random ratio < 2). Finally, using a measure of functional similarity between any pair of areas defined by the degree of their shared inputs or outputs^20^, the more functionally related two areas are, the more likely they are to be connected (Fig. 2d).

**Fig. 2.**
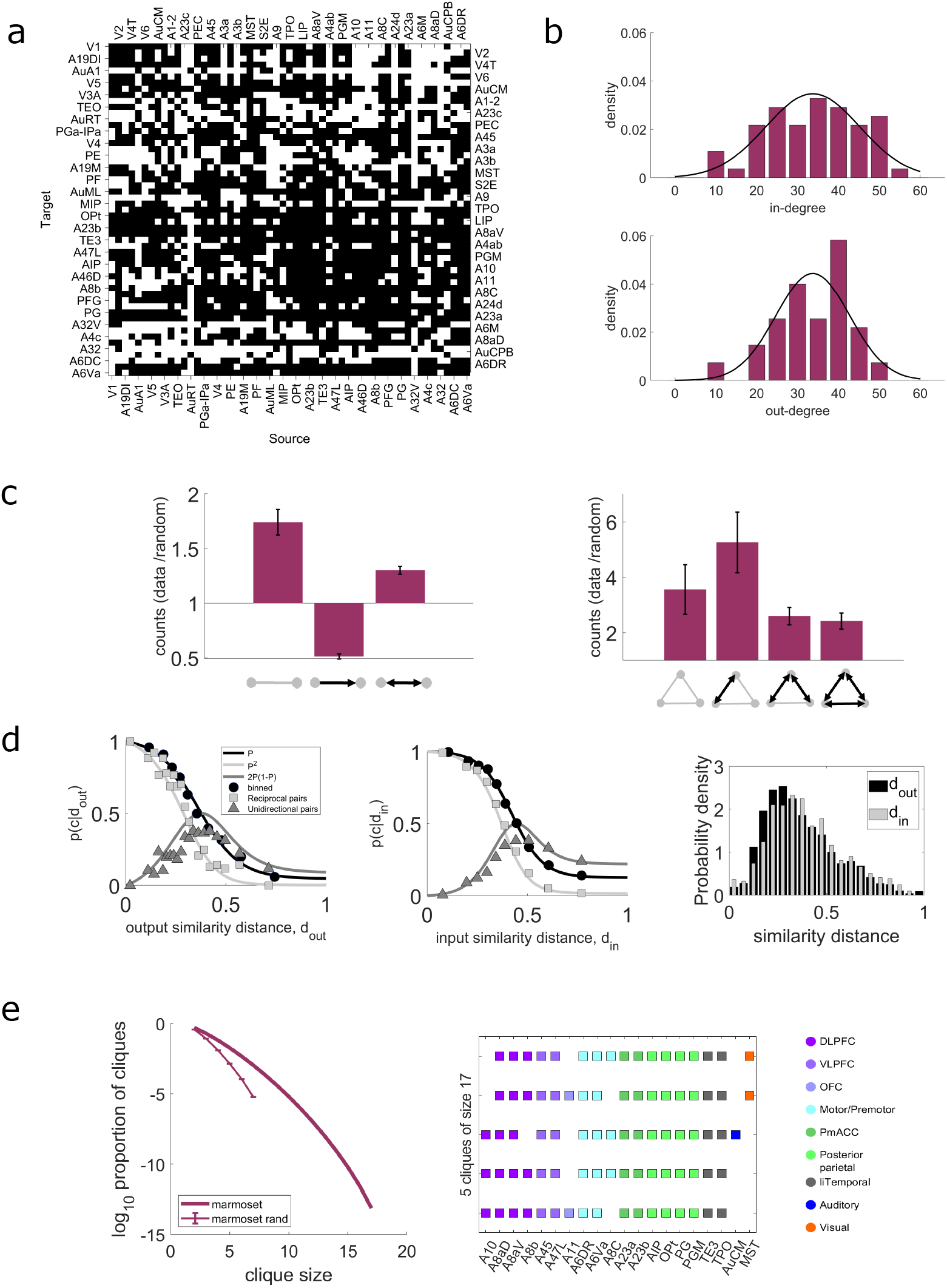
Marmoset network connectivity properties. **(a)** The edge-complete subnetwork, in which all inputs and outputs are known, shows a dense matrix of topological connections. Black: existence, white: absence of a connection. **(b)** In- (left) and out- (right) degree distribution of the target areas. Gray lines are Gaussian fits to the data. **(c)** Average fraction of two- (left) and three- (right) node motif counts of the edge-complete subnetwork to the two- and three-node motif counts, respectively, of a randomized version of the edge-complete network keeping the in- and out-degree the same across 100 realizations. Error bars are one standard deviation of these fractions. **(d)** Proportion of connection as a function of the output (left) and input (middle) similarity distance. Black circles are the number of present connections divided by the number of possible connections in the distance bin. Black line is maximum likelihood fit on the unbinned data. Light grey line is prediction for the reciprocal pairs from the fitted black plot and light gray squares are the proportions of reciprocal pairs in the given bin. Dark grey line is the prediction for the unidirectional pairs and dark grey triangles are the proportions of unidirectional pairs in the given bin. Right: Distribution of the output (black) and input (grey) similarity distances. **(e)** Left: Base 10 logarithm of the proportion of cliques as function of the clique size in the data and the average proportion of cliques in 1000 realizations of a random network of same size where the in- and out-degree sequences are the same as in the data (error bars are one standard deviation). Right: 5 cliques of size 17, combinedly formed by 20 areas that constitute the core of the marmoset cortical connectome.

Another network feature used to characterize the structure of heterogeneous dense networks is cliques, which are subnetworks of fully interconnected areas^13,21,22^. The proportion of cliques of any size in the marmoset connectome is much higher than that of a random network with same in- and out-degree sequences, indicating high specificity (Fig. 2e, left). The largest clique size is 17 and there are five such cliques formed by overall 20 areas (Fig. 2e, right) which define the so-called core of the connectome and has 98.42% density. This is broadly compatible with the core reported in Goulas et al. 2019^22^, with small differences being due to the use of the updated dataset^17^ in the present analysis. The remaining areas constitute the so-called periphery, which form a subnetwork with density 44.29%, and the density of the connections between core and periphery areas is 69.07%. The weights of the connections between areas within the core, and within the periphery, are found to be stronger than those between core- and periphery (Supplementary Fig. 2a). Areas of the putative default mode network (DMN), including those in the posterior parietal cortex (PGM, PG, OPt, AIP), posterior cingulate cortex (A23a, A23b) and dorsolateral prefrontal cortex (A8aD, A6DR) lie in the core structure, but not those in the medial prefrontal cortex (A24d, A32, A32V), reflecting recent studies in the marmoset^23,24^.

### Feedforward projections tend to be stronger than the feedback projections

The structural connectivity of the mammalian cortex is characterized by both the weights of connections between areas and by their laminar organization. A structural hierarchy of the macaque cortex has been defined based on the laminar spatial profile of the connections, according to which ascending (feedforward) pathways originate primarily from the supragranular layers and target layer 4 of the target area; conversely, descending (feedback) pathways originate mostly from the infragranular layers, and target supragranular and infragranular layers^10,18,25,26^. A similar description of information flow has been proposed based on the architectonic type of each area leading to a structural model of the cortex that connects connectivity to evolution, and development^27,28^.

In this framework, the global hierarchical organization can be computed based on the percentage of supragranular neurons involved in the different connections: feedforward connections are formed by high percentages of supragranular neurons in source areas, and feedback connections by low percentages. We calculated the percentage of supragranular neurons (SLN) for a given tracer injection as the number of labeled neurons above layer 4 divided by the total number of all labeled neurons found in the source area (Fig. 3a). When multiple injections were placed in the same area, the SLN was calculated as the weighted average value for the injections that revealed a given connection (Fig. 3b). We found that marmoset cortical connections span the entire range (0-1) of possible SLN values (Fig. 3c). Furthermore, we found that predominantly feedforward (*SLN* > 0.5) connections tended to be stronger than predominantly feedback projections (higher mean FLN by a factor of two; Fig. 3d, similarly in macaque, but less pronounced; Supplementary Fig. 3).

**Fig. 3.**
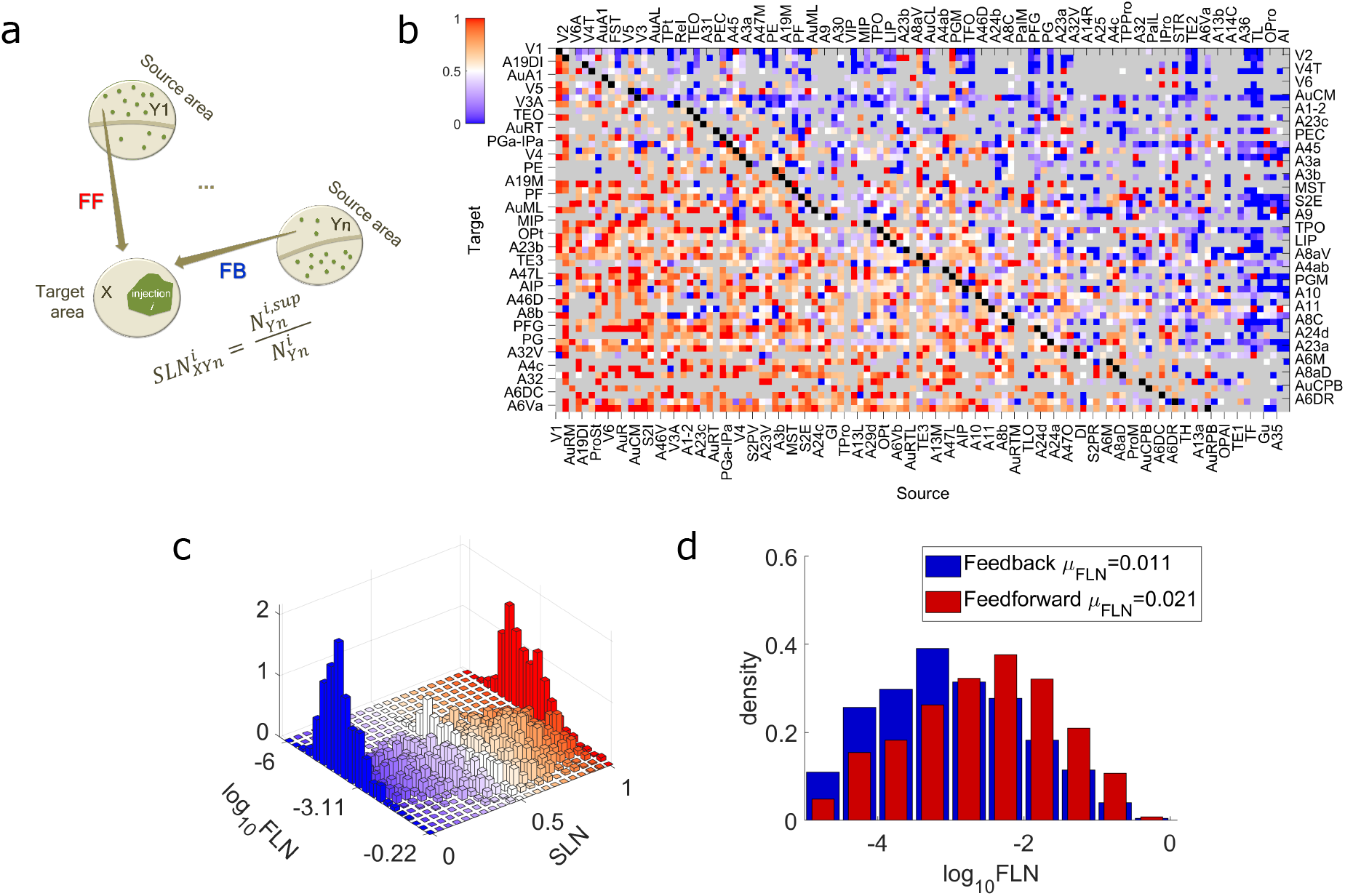
Structural hierarchy. **(a)** Schematic description of the computation of the supragranular labeled neurons found in area *Yn* after the retrograde tracer injection *i* in the cortical area 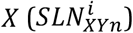. Projections with *SLN* > 0.5 (red entries in (b)) are considered as feedforward projections (FF) and those with *SLN* < 0.5 are feedback projections (FB; blue entries in (b)). **(b)** The SLN matrix. The rows are the 55 target areas and the columns the 116 source areas that provide inputs to each target area, ordered according to the computed hierarchy (Fig. 4a, Supplementary Fig. 4b,right). Each entry in the matrix is the weighted mean supragranular labeled neurons across injections within the same target area. Grey: absence of connections, red: presence of recurrent connections. **(c)** Twodimensional distribution of the FLN and SLN values. The distribution of SLN is not dependent on the strength of connections, except, as expected, at the edges of the distribution formed by very few labeled neurons. **(d)** Distribution of the FLN values of the feedforward connections (red; SLN > 0. 5) and of the feedback connections (blue; SLN < 0.5), with the first being stronger than the latter (higher mean; the two distribution are different (two-sided two-sample Kolmogorov-Smirnov test: *p* = 2.58×10^−40^, Hedges’ *g* effect size: *g* = 0.52), with different mean (two-sided two-sample t-test: *p* = 1.93×10^−43^) but same variance (two-sided two-sample F-test: *p* = 0.86).

### Laminar organization of connections reveals modal hierarchies

To characterize the global hierarchy, we followed a framework used in previous studies^10,26^ in which hierarchical indices are assigned to each area such that for any pair of cortical areas the difference of their hierarchical indices predicts the SLN of their projection (Online Methods). Ordering these indices in Fig. 4a reveals the hierarchy of the edge-complete network. Sensory areas are situated at the bottom of the hierarchy providing feedforward inputs to most other areas, and association areas form mostly feedback projections (Fig. 4a).

**Fig. 4.**
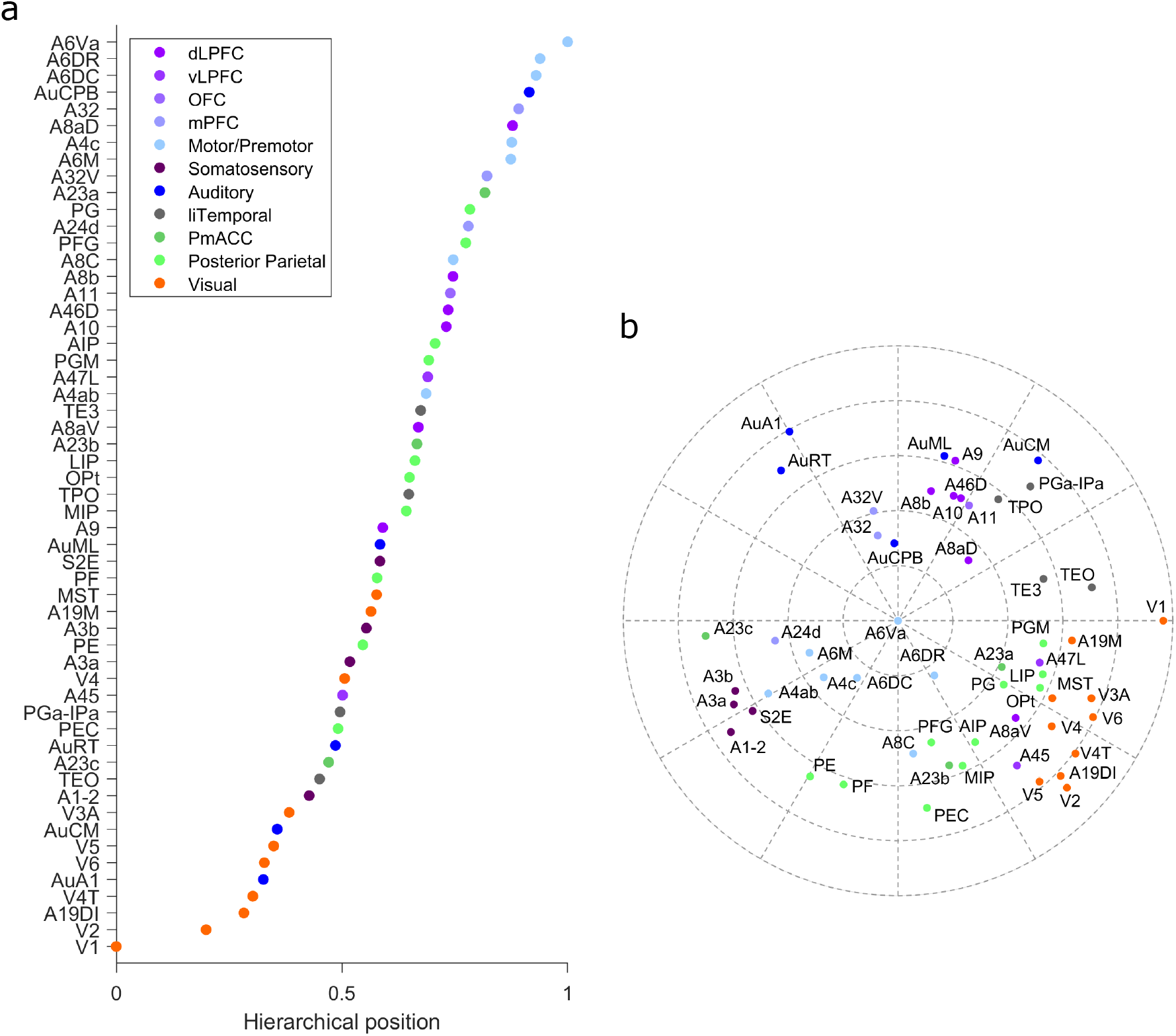
Hierarchical structure. **(a)** Hierarchy of the edge-complete network. **(b)** Two-dimensional representation of the connectivity strength between areas. The radial direction (distance from the outer edge) is defined by the hierarchical position, and the angular distance is given by the inverse of the strength of the connection. It reveals that functionally related areas are grouped together, sensory areas form parallel streams of processing, and different association areas are related to different sensory modalities.

Furthermore, motor areas tend to be concentrated above posterior parietal and prefrontal areas (similarly in macaque cortex but to a lesser extent, Supplementary Fig. 4a), with the ventral premotor cortex being situated at the top of the hierarchy.

This hierarchical ordering aligns the cortical areas based on SLN and doesn’t capture topological properties of the connectivity, e.g. the fact that many connections do not exist. This caveat is mitigated by considering also the core-periphery structure. In this representation, functionally related sensory areas are grouped together in the wings of a “bowtie” graph, with the core occupying the center^29^(Supplementary Fig. 5). The core structure receives effectively stronger feedforward inputs from visual areas, somatosensory and medial prefrontal areas and effectively stronger feedback projections from auditory, posterior parietal and motor areas

The bowtie graph is based on topological binary connections. By including the weights of connections, we can extract information about the overall underlying cortical architecture, represented by a polar plot on a two-dimensional plane^10^. In this representation (Fig. 4b) both the weight of the connections (inverse of the angle between areas) and their hierarchical index (distance from the edge) are considered. Areas deemed to correspond to low hierarchical levels appear in the periphery of the graph, with hierarchical level progressing towards the center. This analysis shows that the marmoset cortical network is better described by a series of modal hierarchies, which converge towards a region formed by multimodal and high-order premotor areas. For example, a hierarchy of visual areas is revealed, grouped together in one quadrant of the plot, which progresses towards parietal areas and frontal areas which are involved in visual cognition (e.g. LIP, Opt, PGM, the frontal eye field [area A8aV] and ventrolateral prefrontal area A47L). The somatosensory and motor areas form another hierarchical grouping in a different quadrant, and areas that are involved in visuomotor integration and planning lie between the visual and somatosensory/ motor clusters (e.g. PEC, MIP and AIP). Finally, the auditory cortex forms a third grouping in a separate quadrant of the plot, with multisensory areas of the temporal lobe (e.g. caudal TPO, PGa-IPa) separating them from visual areas. Interestingly, the prefrontal areas that align best with the auditory hierarchy are those in which single unit activity is related to orientation to sounds in space (e.g. area A8aD) and processing of vocalization sounds (e.g. areas A32/ A32V). Areas associated with the DMN^23,24^ tend to be located near the center of the diagram (e.g. A23a, A6DR).

### Microstructural properties of the cortex reflect the hierarchical organization

An emerging theme of large-scale cortical organization is that biological properties of cortical areas show spatial gradients that correlate with hierarchical level^10,30–32^. Microscale properties are correlated with macroscale connectivity patterns^33^. In addition, previous studies in the macaque^34^ suggested that hierarchical processing is associated with progressively greater numbers of synaptic inputs (leading to greater allocation of space to neuropil, hence lower neuronal densities). The analysis illustrated in Fig. 5 lends support to this hypothesis for the marmoset cortex, in an analysis that combines the newly quantified hierarchical rank and estimate of spine counts on the basal dendritic trees of layer III pyramidal neurons (see Supplementary Table 4 and Supplementary Fig. 6 for sources of data and harmonization of nomenclatures). The average spine counts are highly correlated with hierarchical level (*r* = 0.81). This correlation is equally strong to that against a model based on spatial location of the area along the rostrocaudal axis (Supplementary Fig. 7a,b), which was suggested as another strong predictor of network architecture based on developmental considerations^35^. This constitutes the rostrocaudal axis as proxy for hierarchy.

**Fig. 5.**
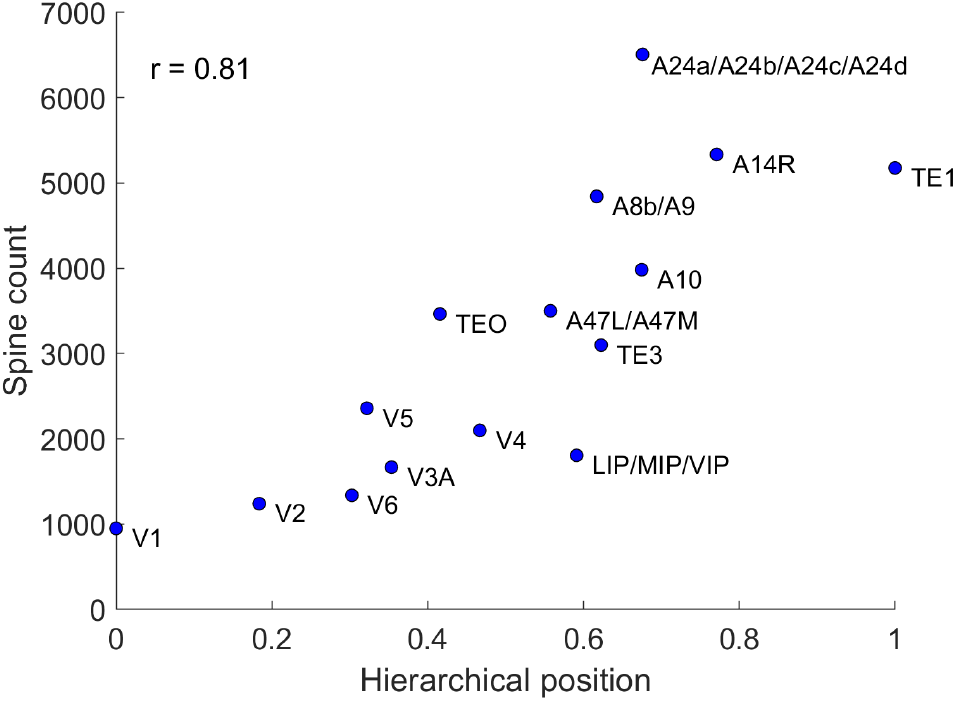
Microstructural properties along the hierarchy. Spine count of basal dendrite in a layer 3 pyramidal neuron is correlated with hierarchical position. *r* is the Pearson correlation.

If we hypothesize that the total number of spines across areas remain constant, we predict that the spine count is inversely proportional to the neural density; along the hierarchy there are fewer bigger neurons with more spines. Based on estimates of neuronal density across the marmoset cortex^36^, we found that neuronal density is indeed decreasing ascending hierarchical levels (Supplementary Fig. 7c, left). However the spine count increases as the inverse cube of neural density suggesting that the total spine count is not constant across areas (Supplementary Fig. 7d). Note of course that the spine count corresponds to the spines of the basal dendrites of the average layer III pyramidal neuron, while the neural density to all neurons. It has been suggested that the number of neurons in a cortical column is a better correlate of hierarchical processing^36^, but we found weaker correlations (Supplementary Fig. 7c, right). Finally, our results support the proposal that both neural density and spine count are good predictors of the laminar origin of projections^37^ since they are highly correlated, and both correlate strongly with the hierarchy.

### Spatial embedding of the connectivity

Parcellated areas in a neocortical network are traditionally considered as nodes of a topological graph, without considering their spatial relationships. However, in addition to its statistical and topological properties, the cortical connectome is spatially embedded. It has been proposed that the metabolic cost of sending information from one area to another increases with distance, being reflected in an exponential distance rule (EDR)^21^ according to which the projection lengths decay exponentially. Incorporation of this attribute in generative models of the connectome helps explain network properties such as efficiency of information transfer, wiring length minimization, 3-motif distribution, and the existence of a core^13,20,21,38^.

To study the spatial organization of the marmoset cortical connectome we used estimates of the distances between the barycenters of cortical areas, obtained with an algorithm that simulates white matter tracts connecting points on the cortical surface^17^. In agreement with observations in other species [macaque^21^, mouse^13^] we found that the wiring distances are normally distributed (Fig. 6a). The distribution of the wiring distances of the pairs for which we have connectivity data (55×116 matrix, Fig. 1c) overlaps with that for all the cortical areas (116×116), indicating that this subset of data is representative of the full network.

**Fig. 6.**
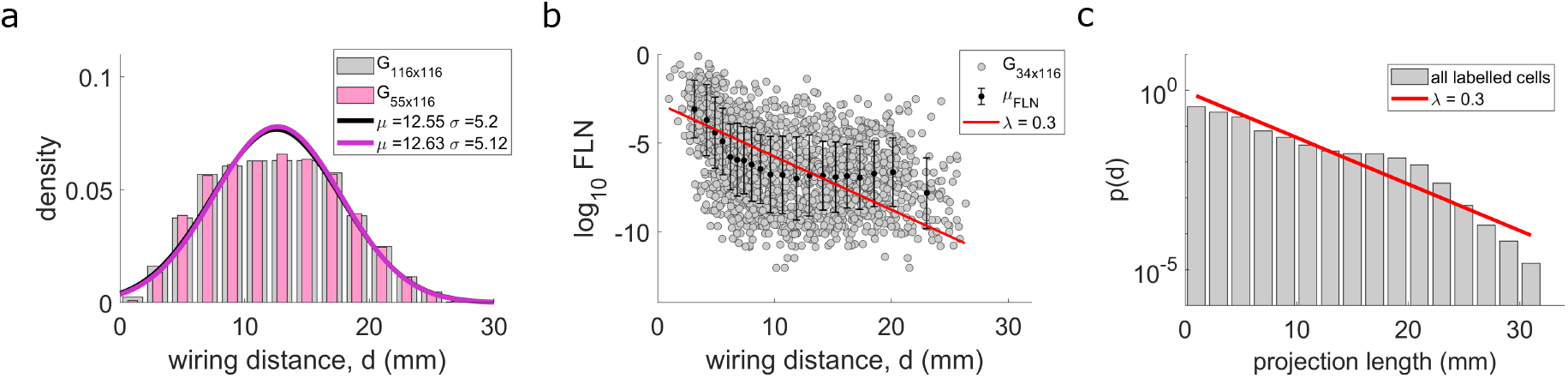
Exponential distance rule. **(a)** Distribution of the inter-areal wiring distances among all 116 areas (gray bars) and between the target-source pairs of areas (pink bars). Bin size is 2 mm. Solid lines are gaussian fits to the data. The normal distributions for these two samples of data indistinguishable (two-sided two-sample Kolmogorov-Smirnov test: *p* = 0.97). **(b)** The base 10 logarithm of the fraction of the extrinsic labeled neurons (*log*_10_*FLN*) as a function of interareal wiring distance. Black dots and error bars are the mean and standard deviation within a window of 173 data points (bin size is 20) and the red plot is the same as in (c). **(c)** The histogram of the projection lengths of all labeled neurons (both intrinsic (551,664 labeled neurons) and extrinsic (1,414,364 labeled neurons), in total 1,966,028 labeled neurons). Bin size is 2 mm and the bars are the counts of the projection lengths lying in the bin size divided by the total number of the projections. The red line is a linear fit to the base 10 logarithm values of the histogram (*log*_10_(*p*(*d*)) = −0.1295 *log*_10_(*projection length*) − 0.0262 giving *p*(*d*) = *ce*^−*λd*^, *λ* ≈ 0.3, *c* ≈ 0.94, where d is the projection length).

We show that the more distant two areas are, the lower the probability of a projection from one to the other, as evidenced by reduced FLN (Fig. 6b). However, analyses based on FLN are anchored on estimates of the borders of cytoarchitectural areas, which in most cases are imprecise^39^. This limitation can be overcome by measuring the wiring distance between each labeled neuron and the corresponding injection site, in an area-independent manner. Based on the stereotaxic coordinates of each injection site and labeled neuron, we calculated the shortest distance across the white matter corresponding to each connection detected in the database (Online Methods and Majka et al. 2020^17^). This included the projection lengths of 1,966,028 labeled neurons, including both those estimated to be in the same area that received the injection (intrinsic connections) and those in other areas. The cost of each neuron to project to longer distances can then be expressed by the distribution of the projection lengths of all the retrogradely labeled neurons (Fig. 6c), which illustrates the probability of a projection length *d*, irrespectively of the areas involved.

As in previous studies in macaque and mouse^13,21,38^ we found that the histogram of axonal projection lengths follows an exponential decay (Fig. 6c) with decay rate *λ* = 0.3; that is, the probability of a projection of length *d* is given by *p*(*d*) = *ce*^−*λd*^. Approximating this with the probability of connections (FLN), it was analytically shown to underlie the log normal distribution of the connectivity weights^21^. In the marmoset this approximation is also valid since the decay of the probability of projection lengths agrees with the *log*_10_*FLN* decay with wiring distance (the red plot falls within the range of the FLN defined by the error bars in Fig. 6b), indicating that the EDR derives the log-normal distribution of the FLN. We should note that the curvature of the average *log*_10_*FLN* and the small bump of projection lengths at distances around 20 mm may suggest the possibility of a more complex relation of the projection lengths distribution. Nevertheless, we showed the projection lengths distribution can be approximated by the EDR, which is an overall statistical property of the cortex.

### Exponential decrease in wiring distance scales with brain volume

Finally, we address the question of how the EDR of cortical connectomes scales across species. It was previously shown that the decay rate of the EDR is larger in macaque than in the mouse following normalization of the distances by the average interareal wiring distance (common template)^13^. This suggests that the larger the brain, the fewer are the long-range connections linking different cortical systems (as shown schematically in Fig. 7, bottom).

**Fig. 7.**
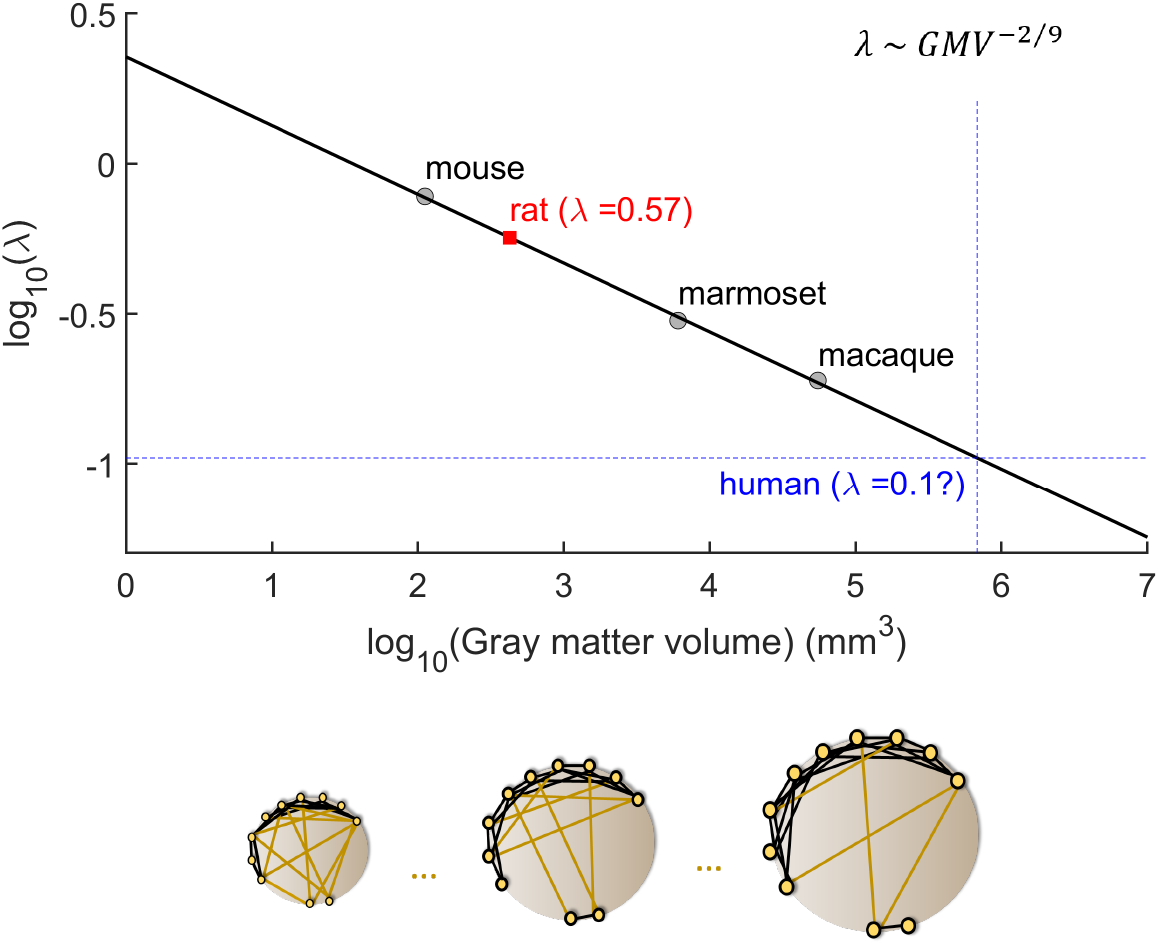
Cortical-connectivity spatial length as a function of brain size: extrapolation to humans. **Top.** The base 10 logarithm of the decay rate of the exponential distance rule (EDR; *λ*) of the mouse, marmoset and macaque, computed in the same way in all three cases. The plot is a linear fit on these three points with a slop of ≈ −2/9 (*log*_10_(*λ*) = −0.2290 *log*_10_(*gray matter volume*) + 0.3559). The red square is the predicted value of the decay rate of the EDR of the rat which is validated by indirect methods^40^ of computing it, and the intersection of the blue dotted lines is the extrapolation of the decay rate of spatial dependence of cortico-cortical connectivity in the human species. **Bottom.** A schematic representation of the decrease of long-range connections as the gray matter gets bigger, showing that the bigger the brain the more local the connectivity.

This principle was upheld in our analysis of the marmoset cortex (Supplementary Fig. 8), where the decay rate in the marmoset shows an intermediate value. Plotting the present data relative to the previous studies in which similar methods were used (macaque and mouse) as a function of the gray matter volume, we found that the decay rate of the EDR (*λ*) scales with grey matter volume following a power law (Fig. 7, top) with an exponent of −2/9. Given that *WM* ~ *GM*^4/3^ (where *WM*: white matter volume and *GM*: gray matter volume)^41^, and if we define the linear dimension *d* as *WM*^1/3^, the decay rate of the axonal projections scales with the inverse square root of the white matter linear dimension, *λ* ~ *d*^-1/2^. It is surprising that the dependence of the characteristic spatial length for EDR is slower than the linear dimension of the white matter, the implication is that the interareal connections become more spatially restricted in a bigger cortex, which presumably is desired for increasing complexity of modular organization.

This power law predicts the decay rate for the rat cortex to be *λ_rat,predicted_* ~ 0.57 *mm*^−1^, which is validated by that estimated indirectly for the rat (*λ_rat,data_* = 0.6 *mm*^−1^) by fitting the EDR to properties of the rat connectome^40^. Finally, using this relation we can extrapolate the decay rate of the projection lengths of the human cortical connectome, which is predicted to be *λ_human, extrapolated_* ~ 0.1 *mm*^−1^ (Fig. 7).

## Discussion

We studied the statistical, architectonic and spatial characteristics of the marmoset cortical mesoscale connectome, based on the largest available dataset for cellular-level connectivity in a primate brain (http://marmosetbrain.org). Our main findings are threefold. First, the marmoset cortical connectome is highly dense at the inter-areal level, characterized by high heterogeneity of inputs and outputs as well as connectivity weights. Moreover, connections are highly specific, evaluated by the presence and absence of reciprocal connections, the dependence of connections on the functional similarity, the distribution of cliques, and the core-periphery structure. Second, based on the laminar origin of the projections we also provided here, for the first time, a quantified hierarchical structure of the marmoset cortex which, in conjunction with connectivity weights, revealed parallel processing streams of sensory and association areas which converge towards a highly connected core. Third, inter-areal connections obey the same exponential distance rule for marmoset as for macaque and mouse cortex. Intriguingly, the characteristic spatial length of the inter-areal connections revealed an allometric scaling rule as a function of the brain size among mammals, leading to a predicted value for the human cortex that can be tested experimentally.

### Scaling across species

Marmosets are New World monkeys, a group which shared a last common ancestor with macaques and humans approximately 43 million years ago^42^. In contrast, the divergence between rodents and primates is estimated to have occurred around 80 million years ago^43^. Here we provide evidence towards the common and species-unique cortical connectivity properties. We show that the connectivity weights of the marmoset are log-normally distributed, similar to that of the macaque^9^ and mouse^1,14^, indicating that this is a general property of the mammalian cortico-cortical connections. In addition, the range of connectivity weights encompasses five orders of magnitude, with a gradual increase in mean connectivity weight as the brain gets smaller (Supplementary Fig. 9). Current estimates indicate ~40 cytoarchitectural areas in the mouse brain, excluding subdivisions of the hippocampal formation^1^, 116 in the marmoset^44^, and 152 in the macaque^45^. Thus, one possibility to account for the above observations is that the dilution of connectivity reflects a gradual redistribution of connections across a larger number of nodes, in larger brains. To test this, we compared the density of the edge-complete graph for available data obtained in the macaque, mouse and marmoset. Perhaps surprisingly, the results (Supplementary Fig. 10) revealed that this is not the case: the density of the cortico-cortical graph is very similar in macaque and marmoset, despite substantial differences in cortex mass and number of cytoarchitectural areas. The above conclusion is robust across application of different thresholds for what is considered a valid connection, an analysis which minimizes the possibility of artifacts related to assignment of cells to adjacent areas, due to uncertainty in histological assessment of borders. Both primates differ from the mouse, in which the graph density is much higher irrespective of the threshold applied (Supplementary Fig. 10). Thus, whereas one may expect an inverse relationship between connection densities and brain volume^14,46^, our results suggest a difference between primates and rodents.

Since FLN can be viewed as area-dependent probability of inter-areal connection, our data suggest scaling of the weights of cortico-cortical connections in such a way that the larger the brain, the larger the proportion of very sparse connections. Extrapolation of these results suggest that the human cortex is likely to be characterized by a comparatively more distributed architecture, showing an even larger proportion of numerically sparse connections. Further, given the scaling of the EDR with brain size, the data also predicts that the larger human brain likely shows a more marked predominance of local connectivity, resulting in a larger number of subnetworks linked by a core (Fig. 7, bottom), as suggested^5^.

### Comparative aspects of the cortical network properties

We have shown that both the marmoset and macaque connectomes exhibit high and similar density. In addition, their in- and out-degree distributions, and the 2- and, 3-node motif distributions, are similar (Supplementary Figs. 1c, d, 11), indicating conserved topological properties independent of the brain size. Another conserved property is that functionally related areas are most likely to be connected. This is related to the wiring distance, with the decay rate of the probability of connections of the marmoset following closely that of the macaque (Supplementary Fig. 12)^20^. Furthermore, the marmoset cortex has more fully interconnected large subnetworks (clique size > 6) in comparison with the macaque (Supplementary Fig. 2b), supporting the view that the smaller the brain, the higher the interconnectivity, even among primates. Finally both the marmoset (Fig. 2e, right) and the macaque^21^ show a similar core structure consisting mainly of association areas.

### Variability of injections

Repeated injections in what are currently considered single cytoarchitectural areas (Supplementary Table 5) revealed variability in their patterns of afferent connections (Supplementary Fig. 13a). In similar studies of the macaque and mouse connectomes^9,14,47^ it has also been shown that the connectivity weights are variable, but less so in the mouse compared to the macaque. One interpretation of these observations is that connectivity patterns across individuals are more consistent in smaller brains, which have fewer subdivisions. This could result from differences in postnatal refinement of patterns of connections in different individuals, which could be more significant in light of more complex behaviors and interactions with the environment. However, other potential sources are within area variability (e.g. differences between the connections of regions serving foveal and peripheral vision^48^, and between parts of the motor cortex related to limb and face movements^49^), hemispheric differences (a subject for which little is known in non-human primates), and those related to the characteristics of individual injections^17^. In previous studies there was deliberate targeting of the same part of the area across subjects^,47^, while in our sample the injections covered different parts across and within subjects (Supplementary Table 2). Thus, the macaque samples were inherently homogeneous, while ours may better reflect the real variability of connections of cytoarchitectural areas. In order to assess this variability thoroughly, a larger statistical sample is required. Nevertheless, the qualitative results based on the weighted values (the FLN and SLN) should be robust to appropriately applied thresholds based on variability across injections. As an example, we show that the decay of the FLN with distance is not affected by considering only the FLN with smaller coefficient of variation across injections in the same target area (Supplementary Fig. 13b).

Another potential source of variability is the inclusion of injections that crossed the estimated borders between two cytoarchitectural areas^17^. This is intrinsically difficult to evaluate in many cases, since the borders of many areas are not sharp^39^. Here injections were assigned to areas based on estimates of how close the injection was to the border and of percentages of the injections contained in each area (see Discussion and Supplementary Table S1 in Majka et al. 2020^17^). However, considering only the injections estimated to be confined at least 80% within the target area (120 injections in 50 target areas; Supplementary Fig. 14b), or injections confined 100% within a target area (79 injections in 34 target areas; Supplementary Fig. 14c) does not substantially affect our conclusions (Supplementary Figs. 10, 15, 16).

### Hierarchical organization

Our analysis of the hierarchical structure of the marmoset cortex indicates that some of the premotor areas, including the ventral premotor cortex (area 6Va), are situated at the top of the hierarchy (Fig. 4). In other words, such areas form a large proportion of projections with characteristics of feedback projections (low SLN). This appears in conflict with the data so far obtained in the macaque, in which association areas such as the prefrontal cortex lie at the highest hierarchical levels^10^ (Supplementary Fig. 4). In addition, the intermediate functional groups were less clearly differentiated. To some extent this may simply reflect differences in the availability of data. For example, to date data of the macaque cortical network does not include injections in ventral premotor cortex (areas F4/F5). Conversely, data obtained in the rostral part of the superior temporal polysensory cortex (area TPO, or “STPr” in the macaque) are not available for the marmoset, where only the caudal part of TPO was injected. Functionally, if the final goal for the cortex is to generate behaviors, it could be expected that the flow of information culminates in motor areas involved in higher-order planning of sequences of movements, such as A6Va and A6DR, which integrate stimulus-initiated and internally-initiated information, towards generation and evaluation of actions. The ventrolateral posterior region of the frontal lobe has expanded considerably in human evolution^15^, including the emergence of Broca’s area in the human brain, suggesting a high-order station for integration of information from various sources, towards generation of complex behavior. However, including the orthogonal dimensions of the presence/absence of connections and weights of connections enabled us to obtain a more comprehensive insight of the inter-areal architecture, where there are parallel sensory streams associated with different higher-level areas. Given the larger number of target areas and pathways studied, this configuration appears clearer than in previous studies of the macaque cortex^10^.

### Future directions: Large scale models of the mammalian brain

One of the principal open problems in systems neuroscience is understanding the structure-to-function relationship. Towards achieving this goal, there is an increasing interest in modeling wholebrain dynamics, as opposed to modeling individual areas. Early models incorporated neuroimaging-based structural connectivity data^50,51^. The major advantage of these studies is that they can model the human brain based on noninvasive data. However, there are important caveats to these modeling studies including the low resolution of the imaging techniques and, critically, unidirectionality of the resulting structural connectivity matrix. On the other hand we show that reciprocity of connections, and absence of connections, are prominent attributes of the connectome (Supplementary Fig. 1) which begs the question whether and how unidirectional connections are important to functions. More recently, there have been a series of large-scale network models that incorporate the weighted and directed structural connectivity obtained via retrograde tracing methods, as well as the hierarchical organization of the areas based on the laminar distribution of the projections^10–12^. The results presented here provide a foundation for a future large-scale network model of the marmoset cortex, which will serve to clarify the computations underlying marmoset brain function and behavior. Ultimately, the increased knowledge of the scaling properties of the cortical cellular network in non-human primates, together with large scale in silico models of other mammals and comparisons of data obtained with neuroimaging techniques^52^, will help bridge the gap between animal models and humans, leading to a better understanding of normal and pathological functions, as well as brain evolution.

## Online methods

### Connectivity data

The marmoset connectivity data consists of the first complete large-scale cortico-cortical connectivity dataset which is available through the Marmoset Brain Connectivity Atlas portal (http://marmosetbrain.org). The detailed methods regarding data collection have been described elsewhere^17,53^. In brief, 143 retrograde tract-tracing experiments were performed in 52 young adult (1.3 – 4.7 years) common marmosets (Callithrix jacchus; 31 male and 21 female), using six types of retrograde tracers (DY (diamidino yellow, 35 injections), FR (fluororuby: dextran-conjugated Tetramethylrhodamine, 35 injections), FB (Fast blue, 29 injections), FE (fluoroemerald: dextran-conjugated fluorescein, 23 injections), CTBgr and CTBr (cholera toxin subunit B, conjugated with Alexa 488 (12 injections) or Alexa 594 (9 injections) respectively)). These 143 injections were made in 55 cortical areas, some of which received more than one injection (Supplementary Tables 1,2,5). All experiments conformed to the Australian Code of Practice for the Care and Use of Animals for Scientific Purposes and were approved by the Monash University Animal Experimentation Ethics Committee^17^. The use of retrograde tracers allowed unambiguous, quantized visualization of individual cell bodies and their precise location relative to cortical layers, which subsequently allowed for precise counting of the labeled cells. Each injection of a retrograde tracer in a cortical area (named as *target area*) results in labelling the neurons that project to it. It has been shown that the majority of the projections to the injected site stem from within the same cortical area^9,47^. Similarly, in the marmoset most of the projections are from within the injected area, but here we don’t consider these connections. Based on the parcellation under consideration^44^ the labeled neurons found in each cortical area (referred to as *source areas*) are counted and categorized based on their laminar position. If the labeled neurons lie above layer IV they are categorized as supragranular labeled neurons and infragranular neurons otherwise. At the same time their stereotaxic coordinates are measured to allow area-independent analyses.

By normalizing the number of labeled neurons in each cortical area (other than the target area) with the total number of labeled neurons in all cortical areas (except the target area) in the same hemisphere, we obtained the so called *fraction of extrinsic labeled neurons* (*FLN*), which are thought to represent the connection weight from the source area to the target area^26^. Specifically, if *X* is an injected cortical area with a retrograde fluorescent tracer, then the fraction of labeled neurons found extrinsic to it, for example in area *Y*, is defined as 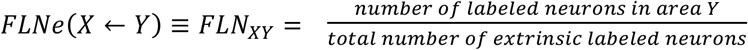. The *FLN_XY_* can be interpreted as the probability of an extrinsic labeled neuron that projects into the target area *X*, is in area *Y*. In Figs. 1b,c the arithmetic average value of the FLN for each target-source pair across injections within the same target area is shown. The bars in the density plot in Fig. 1d (as well as in all density plots accordingly) are the counts of *log*_10_*FLN* values falling in each bin, divided by the bin size (bin size = 0.5) and by the total number of the non-zero FLN values (3474 out of 55×116 = 6380 in total possible interareal connections were present). Within an area the FLN value is set to zero, and therefore also excluded from the density plot. The line is the maximum likelihood Gaussian fit on the *log*_10_*FLN* values.

### Network-related properties

For the topological properties of the connectome we binarized the FLN connectivity matrix (Fig. 1c) by assigning the value 1 (presence of connection) when *FLN* > 0 and 0 (absence of connection) otherwise and considered the edge-complete *N×N* (*N* = 55) network where all inputs and outputs are known (Fig. 2a). The *in-degree* of an area 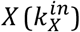 is the number of inputs to this area, meaning the number of areas that project to it. The *out-degree* of an area 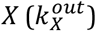 is the number of outputs from this area, meaning the number of areas that this are projects to. In Fig. 2b the density of in- and out-degree of the edge-complete subnetwork (excluding self-connections) binned with bin size 5. The height of each bar denotes the counts divided by the bin size and the total number of areas (*N*). The gray lines are the maximum likelihood Gaussian fits on the normalized in- and out-degree values. The clique size *k* is a *k×k* subnetwork that is fully connected (100% density). In Fig. 2e, left, and Supplementary Fig. 2b for a clique of size k we plot the base 10 logarithm of cliques found in the edge-complete network divided by the maximum number of cliques of size *k* that could be found in the edge-complete network, by taking the *n* choose *k* combinations. We plot the same also for the average random network of same size with same in- and out-degree sequences. The probability of connections as a function of similarity distance (Fig. 2d) was computed following the same method as in Song et al. 2014^20^.

### Hierarchical structure

The fraction of labeled neurons found in the supragranular layers of the source area can be used to calculate the hierarchical rank of each area and it is related to hierarchical distance^10,25,26^. This fraction of labeled supragranular layer neurons (*SLN*) is given by 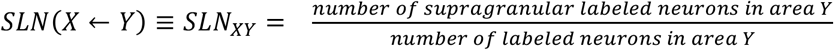, where *X* is the area injected with retrograde tracer (target area) and *Y* is the source area whose neurons project to area *X*. In Fig. 3b the weighted average across injections in the same target area is shown. The areas are ordered with increasing hierarchical index values (Supplementary Fig. 4b,right). Areas APir, Pir, Ent and A29a-c are not shown in the matrix because a layer 4 could not be identified and therefore the SLN is not defined. In Fig. 3c the bars are counts of FLN and SLN within the corresponding bin size (bin size of SLN = 0.05, bin size of *log*_10_*FLN* = 0.289) divided by the bin sizes of SLN and FLN and the total number of the non zero SLN values. In Fig. 3d we categorized the FLN values based on whether their corresponding SLN is greater than 0.5 corresponding to a feedforward projection, and less than 0.5 to a feedback projection, and plotted the probability density as in Fig. 1d.

The hierarchical index for each cortical area *h_i_* (Fig. 4a) is computed via a beta-regression model^54^, where for any target - source pair of areas the difference of their indices can predict the SLN in the source area, as was done for the macaque cortical areas^10,26^. This relationship is expressed through the following equation: *SLN*(*X* ← *Y*) ≈ *g*^−1^(*h_X_*− *h_Y_*), where *g*^−1^ is the logit link function. To obtain the hierarchical indices we used the model fitting function “betareg” in R software, which results in high correlation between predicted and observed SLN values (Supplementary Fig. 17c). Nevertheless, a linear regression model, as in the case of the macaque hierarchy gives similar results (Supplementary Fig. 17a,b). In the model we considered the SLN values of all existing projections from all the injections.

The circular embedding in Fig. 4b is a polar plot of the target areas *A_i_*, with radial coordinate 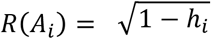 and angular coordinate *θ*(*A_i_*) = *θ_i_*, where *θ_i_* is the angle assigned to each area such that −*log*_10_(*FLN*(*A_i_, A_j_*)) = *r* min(|*θ_i_* − *θ_j_*|, 2*π* − |*θ_i_* − *θ_j_*|), where r is a free parameter, and computed following the same method applied to the macaque cortical areas^10^. The angle of V1 area was assigned to be zero and the system of coordinates were shifted such that the highest area in the hierarchy is at the center of the plot.

### Wiring distances and EDR

If *X* and *Y* are two cortical areas, then the wiring distance *d* ≡ *d_X↔Y_* between them is defined as the shortest path through the white matter, avoiding the grey matter, between their barycenters. The definition is the same as in the studies were the wiring distance of the macaque and mouse was measured and used for the EDR^13,21^. The details of the way the wiring distance were computed can be found in the resource paper of the connectivity data^17^. In brief, the shortest path between the barycenters of two areas was computed by simulating three dimensional trajectories between the areas, where each voxel in the three-dimensional template of the marmoset cortex was assigned different viscosity parameters. The fastest trajectory corresponded to the shortest path. The interareal wiring distance were used in Figs. 6a,b and Supplementary Fig. 12b. For the EDR (Fig. 6c and Supplementary Fig. 8) we used the projection lengths of each labeled neuron, from the injection site to its coordinates (after projecting them to the midthickness surface in order to avoid bias between distances of supragranular and infragranular neurons) measured with the same method as described above. In the EDR plots, each bar represents the counts of the projection lengths lying on the bin divided by the total number of projection lengths (1,966,028 in total) including the projections lengths of the labeled neurons found within the injected area. The red plot in Figs. 6b,c is the linear fit to the log bar plot of the projection lengths, as applied in previous studies^13,21^. Similar fits are also drawn in the common template case (Supplementary Fig. 8).

### Local microstructural properties

We extracted the spine count of marmoset cortical areas from studies where the same method was used (intracellular injection of lucifer yellow), and the same type of spines have been measured (at the basal dendrites of the average pyramidal neuron in layer III) in marmosets of the same age as the ones of the current study (from 18 months to 4.5 years old). We have collected the spine count for 15 cortical areas based on the nomenclature of the papers, which correspond to 22 cortical areas according to the Paxinos et al. 2012^44^ parcellation. Details of the spine count and the corresponding references are shown in Supplementary Table 4 and Supplementary Fig. 6. In Fig. 5, the hierarchical values of the 22 areas have been normalized to 1 and then averaged among areas that correspond to the same spine count (for example the hierarchical index of the area A8b/A9 is the average normalized hierarchical index of areas A8b and A9). In Supplementary Fig. 6b we show that if we instead keep the hierarchical rank of each area and duplicate the spine count for the merged areas (for example area A8b has the same spine count with area A9 but different hierarchical index) the correlation of the spine count is still high. The brain volumes of the marmoset, macaque, mouse, rat and human have been obtained from the literature^41^ (Supplementary Table 6).

### Data availability

The cortico-cortical connectivity datasets analyzed in the current study are available under the terms of Creative Commons Attribution-ShareAlike 4.0 License and publicly available through the Marmoset Brain Connectivity Atlas portal (http://marmosetbrain.org).

### Code availability

Software was written in the MATLAB (https://www.mathworks.com/products/matlab.html), R (https://www.r-project.org/) and Python (https://www.python.org/) programming languages, based on the algorithms of the corresponding published articles and are available upon reasonable request.

## Author contributions

Conceptualization and supervision: XJW, MGPR, DKW. Methodology: PTh, XJW, MGPR, PM, Data Analysis: PTh, PM, Data Curation: MGPR, PM, DHR, PTh. Writing: PTh, MGPR, XJW. Review & Editing: all authors.

## Acknowledgements

The authors would like to thank Henry Kennedy, Kenneth Knoblauch, Maria Ercsey-Ravasz for discussions and Jorge Jaramillo for feedback on the manuscript. Funding was provided by the ONR Grant N00014-17-1-2041, US National Institutes of Health (NIH) grant 062349, and the Simons Collaboration on the Global Brain program grant 543057SPI to XJW; the Australian Research Council grant DP140101968, CE140100007 to MR; the International Neuroinformatics Coordinating Facility and Seed Funding Grant to PM.

## References

1. Oh, S. W. et al. A mesoscale connectome of the mouse brain. Nature 508, 207–214 (2014).

2. Knox, J. E. et al. High-resolution data-driven model of the mouse connectome. Netw. Neurosci. 3, 217–236 (2019).

3. Harris, J. A. et al. Hierarchical organization of cortical and thalamic connectivity. Nature 575, 195–202 (2019).

4. Glasser, M. F. et al. The Human Connectome Project’s neuroimaging approach. Nat. Neurosci. 19, 1175–1187 (2016).

5. Buckner, R. L. & Krienen, F. M. The evolution of distributed association networks in the human brain. Trends Cogn. Sci. 17, 648–665 (2013).

6. Solomon, S. G. & Rosa, M. G. P. A simpler primate brain: the visual system of the marmoset monkey. Front. Neural Circuits 8, 1–24 (2014).

7. Miller, C. T. et al. Marmosets: a neuroscientific model of human social behavior. Neuron 90, 219–233 (2016).

8. Bakola, S., Burman, K. J. & Rosa, M. G. P. The cortical motor system of the marmoset monkey (Callithrix jacchus). Neurosci. Res. 93, 72–81 (2015).

9. Markov, N. T. et al. A weighted and directed interareal connectivity matrix for macaque cerebral cortex. Cereb. Cortex 24, 17–36 (2014).

10. Chaudhuri, R., Knoblauch, K., Gariel, M.-A., Kennedy, H. & Wang, X.-J. A large-scale circuit mechanism for hierarchical dynamical processing in the primate cortex. Neuron 88, 419–431 (2015).

11. Mejias, J. F., Murray, J. D., Kennedy, H. & Wang, X.-J. Feedforward and feedback frequencydependent interactions in a large-scale laminar network of the primate cortex. Sci. Adv. 2, e1601335 (2016).

12. Joglekar, M. R., Mejias, J. F., Yang, G. R. & Wang, X.-J. Inter-areal balanced amplification enhances signal propagation in a large-scale circuit model of the primate cortex. Neuron 98, 222–234. e8 (2018).

13. Horvát, S. et al. Spatial embedding and wiring cost constrain the functional layout of the cortical network of rodents and primates. PLoS Biol. 14, e1002512 (2016).

14. Gămănuţ, R. et al. The mouse cortical connectome, characterized by an ultra-dense cortical graph, maintains specificity by distinct connectivity profiles. Neuron 97, 698–715.e10 (2018).

15. Chaplin, T. A., Yu, H.-H., Soares, J. G. M., Gattass, R. & Rosa, M. G. P. A conserved pattern of differential expansion of cortical areas in simian primates. J. Neurosci. 33, 15120–15125 (2013).

16. Sasaki, E. Prospects for genetically modified non-human primate models, including the common marmoset. Neurosci. Res. 93, 110–115 (2015).

17. Majka, P. et al. Open access resource for cellular-resolution analyses of corticocortical connectivity in the marmoset monkey. Nature Communications 11, 1–14 (2020).

18. Felleman, D. J. & Van Essen, D. C. Distributed hierarchical processing in the primate cerebral cortex. Cereb. Cortex 1, 1–47 (1991).

19. Milo, R. et al. Network motifs: simple building blocks of complex networks. Science 298, 824–827 (2002).

20. Song, H. F., Kennedy, H. & Wang, X.-J. Spatial embedding of structural similarity in the cerebral cortex. PNAS 111, 16580–16585 (2014).

21. Ercsey-Ravasz, M. et al. A predictive network model of cerebral cortical connectivity based on a distance rule. Neuron 80, 184–197 (2013).

22. Goulas, A., Majka, P., Rosa, M. G. P. & Hilgetag, C. C. A blueprint of mammalian cortical connectomes. PLoS Biol. 17, e2005346 (2019).

23. Buckner, R. L. & Margulies, D. S. Macroscale cortical organization and a default-like apex transmodal network in the marmoset monkey. Nat. Commun. 10, 1–12 (2019).

24. Liu, C. et al. Anatomical and functional investigation of the marmoset default mode network. Nat. Commun. 10, 1–8 (2019).

25. Barone, P., Batardiere, A., Knoblauch, K. & Kennedy, H. Laminar distribution of neurons in extrastriate areas projecting to visual areas V1 and V4 correlates with the hierarchical rank and indicates the operation of a distance rule. J. Neurosci. 20, 3263–3281 (2000).

26. Markov, N. T. et al. Anatomy of hierarchy: feedforward and feedback pathways in macaque visual cortex: cortical counterstreams. J. Comp. Neurol. 522, 225–259 (2014).

27. Barbas, H. General cortical and special prefrontal connections: principles from structure to function. Ann. Rev. Neurosci. 38, 269–289 (2015).

28. García-Cabezas, M. Á., Zikopoulos, B. & Barbas, H. The Structural Model: a theory linking connections, plasticity, pathology, development and evolution of the cerebral cortex. Brain Struct. Funct. 224, 985–1008 (2019).

29. Markov, N. T. et al. Cortical high-density counterstream architectures. Science 342, 1238406–1238406 (2013).

30. Burt, J. B. et al. Hierarchy of transcriptomic specialization across human cortex captured by structural neuroimaging topography. Nat. Neurosci. 21, 1251–1259 (2018).

31. Fulcher, B. D., Murray, J. D., Zerbi, V. & Wang, X.-J. Multimodal gradients across mouse cortex. PNAS 116, 4689–4695 (2019).

32. Wang, X.-J. Macroscopic gradients of synaptic excitation and inhibition in the neocortex. Nature Reviews Neuroscience 21, 169–178 (2020).

33. Scholtens, L. H., Schmidt, R., de Reus, M. A. & van den Heuvel, M. P. Linking macroscale graph analytical organization to microscale neuroarchitectonics in the macaque connectome. J. Neurosci. 34, 12192–12205 (2014).

34. Elston, G. N. & Rosa, M. G. Morphological variation of layer III pyramidal neurones in the occipitotemporal pathway of the macaque monkey visual cortex. Cereb. Cortex 8, 278–294 (1998).

35. Cahalane, D. J., Charvet, C. J. & Finlay, B. L. Systematic, balancing gradients in neuron density and number across the primate isocortex. Front. Neuroanat. 6, 1–12 (2012).

36. Atapour, N. et al. Neuronal distribution across the cerebral cortex of the marmoset monkey (callithrix jacchus). Cereb. Cortex 29, 3836–3863 (2019).

37. Beul, S. F. & Hilgetag, C. C. Neuron density fundamentally relates to architecture and connectivity of the primate cerebral cortex. NeuroImage 189, 777–792 (2019).

38. Wang, X.-J. & Kennedy, H. Brain structure and dynamics across scales: in search of rules. Curr. Opin. Neurobiol. 37, 92–98 (2016).

39. Rosa, M. G. P. & Tweedale, R. Brain maps, great and small: lessons from comparative studies of primate visual cortical organization. Philos. Trans. R. Soc. Lond., B, Biol. Sci. 360, 665–691 (2005).

40. Noori, H. R. et al. A multiscale cerebral neurochemical connectome of the rat brain. PLoS Biol. 15, e2002612 (2017).

41. Zhang, K. & Sejnowski, T. J. A universal scaling law between gray matter and white matter of cerebral cortex. PNAS 97, 5621–5626 (2000).

42. Perelman, P. et al. A molecular phylogeny of living primates. PLoS Genet. 7, e1001342 (2011).

43. Foley, N. M., Springer, M. S. & Teeling, E. C. Mammal madness: is the mammal tree of life not yet resolved? Philos. Trans. R. Soc. Lond., B, Biol. Sci. 371, 20150140 (2016).

44. Paxinos, G., Watson, C., Petrides, M., Rosa, M. & Tokuno, H. The marmoset brain in stereotaxic coordinates. (Academic Press, 2012).

45. Saleem, K. S. & Logothetis, N. K. A combined MRI and histology atlas of the rhesus monkey brain in stereotaxic coordinates. (Academic Press, 2012).

46. Ringo, J. L. Neuronal interconnection as a function of brain size. Brain Behav. Evol. 38, 1–6 (1991).

47. Markov, N. T. et al. Weight consistency specifies regularities of macaque cortical networks. Cereb. Cortex 21, 1254–1272 (2011).

48. Palmer, S. M. & Rosa, M. G. P. A distinct anatomical network of cortical areas for analysis of motion in far peripheral vision. Eur. J. Neurosci. 24, 2389–2405 (2006).

49. Burman, K. J., Bakola, S., Richardson, K. E., Reser, D. H. & Rosa, M. G. P. Patterns of afferent input to the caudal and rostral areas of the dorsal premotor cortex (6DC and 6DR) in the marmoset monkey. J. Comp. Neurol. 522, 3683–3716 (2014).

50. Cabral, J., Kringelbach, M. L. & Deco, G. Functional connectivity dynamically evolves on multiple time-scales over a static structural connectome: Models and mechanisms. NeuroImage 160, 84–96 (2017).

51. Demirtaş, M. et al. Hierarchical heterogeneity across human cortex shapes large-scale neural dynamics. Neuron 101, 1181–1194 (2019).

52. Hori, Y. et al. Comparison of resting-state functional connectivity in marmosets with tracer-based cellular connectivity. NeuroImage 204, 116241 (2020).

53. Majka, P. et al. Towards a comprehensive atlas of cortical connections in a primate brain: Mapping tracer injection studies of the common marmoset into a reference digital template. J. Comp. Neurol. 524, 2161–2181 (2016).

54. Cribari-Neto, F. & Zeileis, A. Beta Regression in R. J. Stat. Softw. 34, 1–24 (2010).

