## Supplementary Information for "Structural attributes and principles of the neocortical connectome in the marmoset monkey"

### Supplementary Tables

| Case | Age | Gender | # | Tracer | Target area | Hemisphere |
|---|---|---|---|---|---|---|
| CJ19 | 2y, 24d | M | 1 | FE | V3A | L |
| | | | 2 | FR | V2 | L |
| CJ21 | 3y, 2m, 25d | F | 3 | FB | V3A | L |
| | | | 4 | DY | A19DI | L |
| CJ36 | 2y, 5m, 3d | M | 5 | FR | V2 | L |
| | | | 6 | FB | V6 | L |
| CJ50 | 2y, 11m, 23d | M | 7 | DY | PG | L |
| | | | 8 | FR | V4 | L |
| CJ51 | 3y, 8d | M | 9 | DY | MST | L |
| CJ52 | 2y, 4m, 11d | M | 10 | DY | PEC | L |
| | | | 11 | FR | V6 | L |
| | | | 12 | FE | V6 | L |
| CJ55 | 3y, 10m, 11d | F | 13 | DY | PG | L |
| | | | 14 | FR | LIP | L |
| | | | 15 | FB | OPt | L |
| | | | 16 | FE | MIP | L |
| CJ56 | 1y, 6m, 25d | M | 17 | FR | V5 | L |
| | | | 18 | FB | V5 | L |
| | | | 19 | DY | V4T | L |
| | | | 20 | FE | V5 | L |
| CJ64 | 1y, 5m, 7d | M | 21 | FE | AuML | R |
| | | | 22 | FR | AuCM | R |
| | | | 23 | FB | V4T | R |
| CJ70 | 1y, 4m, 7d | M | 24 | DY | A10 | R |
| | | | 25 | FB | A9 | R |
| | | | 26 | FR | A8aD | R |
| CJ71 | 1y, 6m, 8d | M | 27 | DY | A10 | R |
| | | | 28 | FB | A10 | R |
| | | | 29 | FR | A47L | R |
| CJ73 | 1y, 5m, 12d | M | 30 | FR | A10 | R |
| | | | 31 | FB | A47L | R |
| | | | 32 | DY | A8b | R |
| | | | 33 | FE | A47L | R |
| CJ74 | 1y, 6m, 22d | M | 34 | FR | A8aV | R |
| | | | 35 | FB | A8b | R |
| | | | 36 | DY | A8b | R |
| Cj75 | 1y, 4m, 23d | M | 37 | DY | A8aV | L |
| | | | 38 | FR | AuA1 | L |
| CJ76 | 1y, 6m, 25d | M | 39 | FE | PFG | L |
| | | | 40 | DY | PFG | L |
| | | | 41 | FR | PG | L |
| | | | 42 | FB | LIP | L |
| CJ77 | 1y, 5m, 8d | M | 43 | DY | PF | R |
| | | | 44 | FR | PG | R |
| | | | 45 | FB | OPt | R |
| CJ78 | 2y, 19d | M | 46 | FE | A4ab | L |
| | | | 47 | FB | A4ab | L |
| | | | 48 | DY | A3a | L |
| CJ80 | 1y, 8m, 5d | M | 49 | FB | A23b | R |
| | | | 50 | FR | A23a | R |
| CJ80 | 1y, 8m, 5d | M | 51 | DY | PGM | R |
| CJ81 | 2y, 4m, 19d | F | 52 | FR | V2 | L |
| | | | 53 | FB | V2 | L |
| | | | 54 | DY | V2 | L |
| | | | 55 | FE | V2 | L |
| CJ82 | 2y, 4m, 27d | F | 56 | DY | V2 | L |
| | | | 57 | FR | V1 | L |
| | | | 58 | FE | V1 | L |
| CJ83 | 2y, 5m, 27d | F | 59 | DY | A8b | L |
| CJ84 | 4y, 7m, 5d | F | 60 | FE | A23a | L |
| | | | 61 | FB | PGM | L |
| | | | 62 | FR | A19M | L |
| CJ90 | 3y, 1m | F | 63 | FR | V2 | R |
| | | | 64 | DY | V2 | R |
| CJ94 | 3y, 3m, 11d | F | 65 | DY | A8aV | R |
| | | | 66 | FR | A6DR | R |
| | | | 67 | FB | A6M | R |
| CJ100 | 3y, 7m, 16d | F | 68 | FR | A6DR | R |
| | | | 69 | FE | A6Va | R |
| CJ102 | 4y, 12d | F | 70 | DY | A3b | L |
| CJ108 | 3y, 1m, 13d | M | 71 | FE | A8aV | R |
| | | | 72 | FR | A8aD | R |
| CJ110 | 3y, 1m, 6d | M | 73 | FE | A6DR | L |
| | | | 74 | FR | A6Va | L |
| CJ111 | 3y, 1m, 13d | M | 75 | FE | A6Va | L |
| | | | 76 | FR | A4ab | L |
| CJ112 | 3y, 1m, 18d | M | 77 | FE | A6DC | L |
| | | | 78 | FR | A6Va | L |
| CJ113 | 3y, 2m, 15d | M | 79 | FE | A6M | L |
| | | | 80 | FR | A4ab | L |
| CJ114 | 3y, 4m | M | 81 | FR | A4ab | R |
| | | | 82 | FE | A3b | R |
| CJ115 | 2y, 11m, 16d | F | 83 | FE | A6DC | R |
| | | | 84 | FR | A4c | R |
| CJ116 | 3y, 2m, 12d | F | 85 | FR | A6DR | R |
| | | | 86 | FE | A6Va | R |
| CJ122 | 2y, 10m, 23d | M | 87 | DY | AuRT | R |
| | | | 88 | FR | AuCPB | R |
| | | | 89 | FB | TPO | R |
| | | | 90 | FE | TPO | R |
| CJ123 | 4y, 8m, 12d | F | 91 | FE | A8C | R |
| | | | 92 | FR | A6DC | R |
| CJ125 | 4y, 8m, 19d | F | 93 | FR | A8aV | L |
| | | | 94 | FE | A6DR | L |
| CJ146 | 2y, 1m, 11d | M | 95 | DY | A23b | L |
| | | | 96 | FR | A23a | L |
| | | | 97 | FB | A23b | R |
| CJ148 | 1y, 7m, 13d | F | 98 | DY | A32V | L |
| CJ153 | 1y, 7m | F | 99 | FR | A23b | L |
| CJ164 | 1y, 6m, 15d | F | 100 | DY | A8b | L |
| CJ164 | 1y, 6m, 15d | F | 101 | FB | A8b | L |
| | | | 102 | CTBgr | A6M | L |
| CJ167 | 2y, 3m, 6d | F | 103 | CTBgr | A24d | R |
| | | | 104 | FB | A24d | R |
| | | | 105 | FR | A24d | R |
| | | | 106 | DY | A23c | R |
| CJ170 | 2y, 3m, 26d | M | 107 | DY | A6M | R |
| | | | 108 | FB | A3a | R |
| | | | 109 | CTBr | A3b | R |
| | | | 110 | CTBgr | S2E | R |
| CJ173 | 2y, 17d | M | 111 | DY | A4ab | R |
| | | | 112 | FB | A1-2 | R |
| | | | 113 | CTBr | PE | R |
| | | | 114 | CTBgr | AIP | R |
| CJ174 | 2y, 7m, 3d | F | 115 | FB | V1 | R |
| | | | 116 | CTBgr | V1 | R |
| | | | 117 | CTBr | V1 | R |
| | | | 118 | CTBr | A10 | R |
| CJ178 | 2y, 3m, 15d | F | 119 | CTBgr | A10 | R |
| | | | 120 | DY | A32V | R |
| | | | 121 | FB | V1 | R |
| CJ180 | 2y, 8m, 29d | M | 122 | FB | AuRT | R |
| | | | 123 | CTBr | TE3 | R |
| | | | 124 | CTBgr | AuML | R |
| | | | 125 | DY | TE3 | R |
| CJ181 | 2y, 2m, 19d | M | 126 | CTBr | A11 | R |
| | | | 127 | CY | A47L | R |
| | | | 128 | CTBgr | AuCM | R |
| | | | 129 | FB | TEO | R |
| CJ182 | 3y, 2m, 16d | M | 130 | CTBr | PGa-IPa | R |
| | | | 131 | CTBgr | V4 | R |
| | | | 132 | DY | V4 | R |
| CJ800 | 2y, 9m, 6d | M | 133 | DY | A46D | R |
| | | | 134 | FB | A46V | R |
| | | | 135 | CTBr | A8aD | R |
| | | | 136 | CTBgr | A45 | R |
| CJ801 | 3y, 6m | F | 137 | DY | A46D | R |
| | | | 138 | CTBgr | A6DR | R |
| | | | 139 | FB | A8C | R |
| CJ802 | 3y, 8m, 15d | M | 140 | CTBr | S2E | R |
| | | | 141 | CTBgr | AuML | R |
| | | | 142 | DY | AIP | R |
| | | | 143 | FB | PG | R |

**Supplementary Table 1 | Demographics of the injections.** List of the marmosets (case), tracers used, hemisphere (L = left, R = right) of the injection, injected area (target area), age (y = years, m = months, d = days) and gender (M = male, F = female) of the marmosets sacrificed for this, and not only, study.

| marmoset | DY | FR | FB | FE | CTBgr | CTBr |
|---|---|---|---|---|---|---|
| | | | tracer | | | |
| CJ19 | | V2 | | V3A | | |
| CJ21 | A19DI | | V3A | | | |
| CJ36 | | V2 | V6 | | | |
| CJ50 | PG | V4 | | | | |
| CJ51 | MST | | | | | |
| CJ52 | PEC | V6 | | V6 | | |
| Cj55 | PG | LIP | OPt | MIP | | |
| CJ56 | V4T | V5 | V5 | V5 | | |
| CJ64 | | AuCM | V4T | AuML | | |
| CJ70 | A10 | A8aD | A9 | | | |
| CJ71 | A10 | A47L | A10 | | | |
| CJ73 | A8b | A10 | A47L | A47L | | |
| CJ74 | A8b | A8aV | A8b | | | |
| CJ75 | A8aV | AuA1 | | | | |
| CJ76 | PFG | PG | LIP | PFG | | |
| CJ77 | PF | PG | OPt | | | |
| CJ78 | A3a | | A4ab | A4ab | | |
| CJ80 | PGM | A23a | A23b | | | |
| CJ81 | V2 | V2 | V2 | V2 | | |
| CJ82 | V2 | V1 | | V1 | | |
| Cj83 | A8b | | | | | |
| CJ84 | | A19M | PGM | A23a | | |
| CJ90 | V2 | V2 | | | | |
| CJ94 | A8aV | A6DR | A6M | | | |
| CJ100 | | A6DR | | A6Va | | |
| CJ102 | A3b | | | | | |
| CJ108 | | A8aD | | A8aV | | |
| CJ110 | | A6Va | | A6DR | | |
| CJ111 | | A6DC | | A6Va | | |
| CJ112 | | A6Va | | A6DC | | |
| CJ113 | | A4ab | | A6M | | |
| CJ114 | | A4ab | | A3b | | |
| CJ115 | | A4c | | A6DC | | |
| CJ116 | | A6DR | | A6Va | | |
| CJ122 | AuRT | AuCPB | TPO | TPO | | |
| CJ123 | | A6DC | | A8C | | |
| CJ125 | | A8aV | | A6DR | | |
| CJ146 | A23b | A23a | A23b | | | |
| CJ148 | A32V | | | | | |
| CJ153 | | A23b | | | | |
| CJ164 | A8b | | A8b | | A6M | |
| CJ167 | A23c | A24d | A24d | | A24d | |
| CJ170 | A6M | | A3a | | S2E | A3b |
| CJ173 | A4ab | | A1-2 | | AIP | PE |
| CJ174 | | | V1 | | V1 | V1 |
| CJ178 | A32V | | V1 | | A10 | A10 |
| CJ180 | TE3 | | AuRT | | AuML | TE3 |
| CJ181 | A47L | | TEO | | AuCM | A11 |
| CJ182 | V4 | | | | V4 | PGa-IPa |
| CJ800 | A46D | | A47L | | A45 | A8aD |
| CJ801 | A46D | | A8C | | A6DR | |
| CJ802 | AIP | | PG | | AuML | S2E |
| | DY | FR | FB | FE | CTBgr | CTBr |

**Supplementary Table 2 | Location and tracer of each injection.** All 143 injections to the 52 marmosets (rows) with the six retrograde tracers (columns). Each cell shows the cortical area that has been injected to the corresponding marmoset with the corresponding tracer. Areas in bold denote right hemisphere, otherwise left hemisphere. In some marmosets (in shaded pink) there has been more than one injection to the same target area with different tracer, in most cases within the same hemisphere but also across hemispheres (marmoset CJ146).

| | |
|---|---|
| *Dorsolateral prefrontal cortex (6/7 target areas)* | |
| **A10 | area 10 of cortex |
| **A8b | area 8b of cortex |
| **A9 | area 9 of cortex |
| **A46D | area 46 of cortex dorsal part |
| **A8aD | area 8a of cortex dorsal part |
| **A8aV | area 8a of cortex ventral par |
| A46V | area 46 of cortex ventral part |
| *Ventrolateral prefrontal cortex (2/5 target areas)* | |
| **A47L | area 47 (old 12) of cortex lateral part |
| **A45 | area 45 of cortex |
| A47M | area 47 (old 12) of cortex medial part |
| A47O | area 47 (old 12) of cortex orbital part |
| ProM | proisocortical motor region (precentral opercular cortex) |
| *Orbitofrontal cortex (1/8 target area)* | |
| **A11 | area 11 of cortex |
| A13a | area 13a of cortex |
| A13b | area 13b of cortex |
| A13L | area 13 of cortex lateral part |
| A13M | area 13 of cortex medial part |
| OPAl | orbital periallocortex |
| OPro | orbital proisocortex |
| Gu | gustatory cortex |
| *Medial prefrontal cortex (3/9 target areas)* | |
| **A32 | area 32 of cortex |
| **A32V | area 32 of cortex ventral part |
| **A24d | area 24d of cortex |
| A14C | area 14 of cortex caudal part |
| A14R | area 14 of cortex rostral part |
| A25 | area 25 of cortex |
| A24a | area 24a of cortex |
| A24b | area 24b of cortex |
| A24c | area 24c of cortex |
| *Motor and premotor cortex (7/8 target areas)* | |
| **A6DR | area 6 of cortex dorsorostral part |
| **A6DC | area 6 of cortex dorsocaudal part |
| **A4ab | area 4 of cortex parts a and b (primary motor) |
| **A6Va | area 6 of cortex ventral part a |
| **A8C | area 8 of cortex caudal part |
| **A6M | area 6 of cortex medial (supplementary motor) part |
| **A4c | area 4 of cortex part c (primary motor) |
| A6Vb | area 6 of cortex ventral part b |
| *Insular and rostral lateral sulcus cortex (0/7 target areas)* | |
| PaIL | parainsular cortex lateral part |
| PaIM | parainsular cortex medial part |
| AI | agranular insular cortex |
| DI | dysgranular insular cortex |
| GI | granular insular cortex |
| IPro | insular proisocortex |
| TPro | temporal proisocortex |
| *Lateral and inferior temporal cortex (4/10 target areas)* | |
| **TE3 | temporal area TE3 (inferior temporal cortex) |
| **TEO | temporal area TE occipital part |
| **PGa/IPa | Area PGa andIPa (fundus of superior temporal ventral area) |
| **TPO | temporo-parieto-occipital association area (superior temporal polysensory cortex) |
| TPPro | temporopolar proisocortex |
| STR | superior temporal rostral area (cortex) |
| TE1 | temporal area TE1 (inferior temporal cortex) |
| ReI | retroinsular area (cortex) |
| TE2 | temporal area TE2 (inferior temporal cortex) |
| TPt | temporoparietal transitional area |
| *Ventral temporal cortex (0 target áreas)* | |
| APir | amygdalopiriform transition area |
| Pir | piriform transition area |
| Ent | entorhinal cortex |
| A35 | area 35 of cortex |
| A36 | area 36 of cortex |
| TF | temporal area TF |
| TL | temporal area TL |
| TH | temporal area TH |
| TLO | temporal area TL occipital part |
| TFO | temporal area TF occipital part |
| *Posterior cingulate, medial and retrosplenial cortex (3/8 target areas)* | |
| **A23a | area 23a of cortex |
| **A23b | area 23b of cortex |
| **A23c | area 23c of cortex |
| A29a-c | area 29a-c of cortex |
| A29d | area 29d of cortex |
| A30 | area 30 of cortex |
| A23V | area 23 of cortex ventral part |
| ProSt | prostriate area |
| *Posterior parietal cortex (10/13 target areas)* | |
| **AIP | anterior intraparietal area of cortex |
| **LIP | lateral intraparietal area of cortex |
| **MIP | medial intraparietal area of cortex |
| **OPt | occipito-parietal transitional area of cortex |
| **PE | parietal area PE |
| **PEC | parietal area PE caudal part |
| **PF | parietal area PF (cortex) |
| **PFG | parietal area PFG (cortex) |
| **PG | parietal area PG |
| **PGM | parietal area PG medial part (cortex) |
| A31 | area 31 of cortex |
| V6A | visual area 6A (posterior parietal medial area) |
| VIP | ventral intraparietal area of cortex |
| *Somatosensory cortex (3/7 target areas)* | |
| **S2E | secondary somatosensory cortex external part |
| **A3b | area 3b of cortex (somatosensory) |
| **A1-2 | areas 1 and 2 of cortex |
| S2I | secondary somatosensory cortex internal part |
| S2PR | secondary somatosensory cortex parietal rostral area |
| S2PV | secondary somatosensory cortex parietal ventral area |
| A3a | area 3a of cortex (somatosensory) |
| *Auditory cortex (5/12 target areas)* | |
| **AuA1 | auditory cortex primary area |
| **AuCM | auditory cortex caudomedial area |
| **AuCPB | auditory cortex caudal parabelt area |
| **AuML | auditory cortex middle lateral area |
| **AuRT | auditory cortex rostrotemporal |
| AuAL | auditory cortex anterolateral area |
| AuCL | auditory cortex caudolateral area |
| AuR | auditory cortex rostral area\ |
| AuRM | auditory cortex rostromedial area |
| AuRPB | auditory cortex rostral parabelt |
| AuRTL | auditory cortex rostrotemporal lateral area |
| AuRTM | auditory cortex rostrotemporal medial area |
| *Visual cortex (10/12 target areas)* | |
| **A19DI | area 19 of cortex dorsointermediate part |
| **A19M | area 19 of cortex medial part |
| **MST | medial superior temporal area of cortex |
| **V1 | primary visual cortex |
| **V2 | visual area 2 |
| **V3A | visual area 3A (dorsoanterior area) |
| **V4 | visual area 4 (ventrolatereral anterior area) |
| **V4T | visual area 4 transitional part (middle temporal crescent) |
| **V5 | visual area 5 (middle temporal area) |
| **V6 | visual area 6 (dorsomedial area) |
| FST | fundus of superior temporal sulcus area of cortex |
| V3 | visual area 3 (ventrolateral posterior area) |

**Supplementary Table 3 | Abbreviations of the names of cortical areas, grouped in the major cortical subdivisions.** Colors correspond to the color code in figures (Fig. 4, and Supplementary Figs. 4,16,17). Asterisks denote the target areas. Target areas are broadly distributed in the cortex, selected from all the major subdivisions, except the insular and rostral lateral sulcus cortex and the ventral temporal cortex.

| Reference | Area in reference | Area in Paxinos et al. 2012 parcellation | Spine Count | Age |
|---|---|---|---|---|
| Aoi et al. 2013 | V1 | V1 | 950 | 4.5 Y |
| Elston et al. 1999 | V2 | V2 | 1240 | 20 -- 28 M |
| Elston et al. 1999 | DM | V6 | 1338 | 20 -- 28 M |
| Elston et al. 1999 | DA | V3A | 1667 | 20 -- 28 M |
| Elston et al. 1999 | PP | LIP/MIP/VIP | 1806 | 20 -- 28 M |
| Elston et al. 1999 | DL | V4 | 2098 | 20 -- 28 M |
| Elston et al. 1999 | MT | V5 | 2359 | 20 -- 28 M |
| Aoi et al. 2013 | Itd/TE3 | TE3 | 3100 | 4.5 Y |
| Elston et al. 1999 | Itc/TEO | TEO | 3465 | 20 -- 28 M |
| Aoi et al. 2013 | 12L+12M | A47L/A47M | 3500 | 4.5 Y |
| Elston et al. 2001 | 10 | A10 | 3983 | 18 M |
| Sasaki et al. 2015 | 8B/9 | A8b/A9 | 4843 | 2.5 Y |
| Elston et al. 1999 | Itr/TE | TE1 | 5176 | 20 -- 28 M |
| Sasaki et al. 2015 | 14r | A14R | 5334 | 2.5 Y |
| Sasaki et al. 2015 | 24 | A24a/A24b/A24c/A24d | 6507 | 2.5 Y |

**Supplementary Table 4 | Spine count data.** Spine count data (fourth column) collected from the literature (first column[1–4]) for 15 cortical areas based on the parcellation in the corresponding source papers (second column), and 22 areas in the Paxinos et al. 2012[5] parcellation (third column). The data were chosen from the literature so that to correspond to sexually matured marmosets (fifth column) since the spine count shows age dependence[4], as well as to correspond to the age range of the marmosets of this study. Specifically, the spine count of the average pyramidal layer three neuron in area V1, TE3, A47L/A47M has 950, 3100, 3500 spines respectively[3], in area A14R, A8b/A9, A24a/A24b/A24c/A24d has 5334, 4843, 6507 spines respectively[4], in area V2, V6, V3A, LIP/MIP/VIP, V4, V5, TEO, TE1 has 1240, 1338, 1667, 1806, 2098, 2359 spine respectively[1], and in area A10 has 3983[2] The spine count in V1 in an earlier study[1] was reported to be smaller but similar (699 spines for marmoset of age 20-28 months old), so we chose to consider the most recent report[3].

| Target Area | Number of Injections | Target Area | Number of Injections | Target Area | Number of Injections |
|---|---|---|---|---|---|
| V2 | 9 | A3a | 2 | A1-2 | 1 |
| V1 | 6 | A46D | 2 | A11 | 1 |
| A6DR | 6 | AIP | 2 | A19DI | 1 |
| A10 | 6 | AuCM | 2 | A19M | 1 |
| A8b | 6 | AuRT | 2 | A23c | 1 |
| A47L | 5 | PFG | 2 | A32 | 1 |
| A4ab | 5 | PGM | 2 | A32V | 1 |
| A6Va | 5 | S2E | 2 | A45 | 1 |
| PG | 5 | TE3 | 2 | A4c | 1 |
| A8aV | 5 | TPO | 2 | A9 | 1 |
| A23b | 4 | V4T | 2 | AuA1 | 1 |
| A6DC | 4 | A8C | 2 | AuCPB | 1 |
| A6M | 4 | V3A | 2 | MIP | 1 |
| V6 | 3 | LIP | 2 | MST | 1 |
| V5 | 3 | Opt | 2 | PE | 1 |
| V4 | 3 | | | PEC | 1 |
| A23a | 3 | | | PF | 1 |
| AuML | 3 | | | PGa-IPa | 1 |
| A24d | 3 | | | TEO | 1 |
| A8aD | 3 | | | | |
| A3b | 3 | | | | |

**Supplementary Table 5 | Multiple injections within each target area.** Number of injections in each of the 55 target areas. Overall there is one area with 9 injections (V2), 4 areas with 6 injections (V1, A6DR, A10, A8b), 5 areas with 5 injections (A47L, A4ab, A6Va, PG, A8aV), 3 areas with 4 injections (A23b, A6DC, A6M), 8 areas with 3 injections (V6, V5, V4, A23a, AuML, A24d, A8aD, A3b), 15 with 2 injections (A3a, A46D, AIP, AuCM, AuRT, PFG, PGM, S2E, TE3, TPO, V4T, LIP, A8C, OPt, V3A), and 19 areas with one injections (A1-2, A11, A19DI, A19M, A23c, A32, A32V, A45, A4c, A9, AuA1, AuCPB, MIP, MST, PE, PEC, PF, PGa-IPa, TEO).

| Reference | mouse | rat | marmoset | macaque | human |
|---|---|---|---|---|---|
| Hofman 1988 | | | 6,090 | 55,100 | 683,000 |
| Zhang and Sejnowski 2000 | 112 | 425 | | | |

**Supplementary Table 6 | Brain volume data.** Gray matter volume data[6,7] in $mm^3$**.** Specifically, the grey matter volume of the mouse and rat ($GMV_{mouse}$ = 112 $mm^3$, $GMV_{rat}$ = 425 $mm^3$) were obtained from the caption of Figure 2 in Zhang and Sejnowskii 2000[7] and the grey matter volume of the marmoset, macaque and human ($GMV_{marmoset}$ = 6090 $mm^3$, $GMV_{macaque}$ = 55100 $mm^3$, $GMV_{human}$ = 683000 $mm^3$) from Table 1 in Hofman 1988[6]. A recent study[8] have measured the volume of the primate neocortex (marmoset: 4,451.98 $mm^3$, macaque: 44,557.89 $mm^3$, no human data) with magnetic resonance. These values are not much different from those shown in the table and when taken into account they don't alter the results shown in Fig. 7 (not shown here). The values shown in the table have been used in Zang and Sejnowski 2000[7] where they showed the power law scaling of the gray matter with white matter volume based on data from many species and we chose to consider these for consistency.

| Area | Rostrocaudal coordinate | Area | Rostrocaudal coordinate | Area | Rostrocaudal coordinate |
|---|---|---|---|---|---|
| A10 | -18.9 | A24d | -10.3 | A23a | -4.3 |
| A9 | -17.2 | STR | -10.3 | PE | -4.3 |
| A46V | -17.1 | AI | -10.2 | PFG | -4.3 |
| A46D | -17.1 | APir | -10.0 | TF | -4.0 |
| A47L | -17.0 | PaIL | -9.9 | A29d | -4.0 |
| A11 | -16.7 | AuRTL | -9.9 | MST | -3.9 |
| A32 | -16.3 | DI | -9.7 | TE3 | -3.9 |
| A47M | -16.3 | AuRT | -9.7 | A30 | -3.9 |
| A13b | -16.2 | A4ab | -9.6 | A31 | -3.7 |
| A32V | -16.0 | AuRPB | -9.5 | A23b | -3.4 |
| A14R | -15.4 | AuRTM | -9.2 | AIP | -3.3 |
| A8aD | -15.2 | GI | -9.1 | TL | -2.8 |
| A8b | -15.2 | A3a | -9.1 | TH | -2.7 |
| A8aV | -15.0 | TE1 | -8.9 | FST | -2.5 |
| A45 | -14.7 | S2PV | -8.7 | PG | -2.3 |
| A13a | -14.5 | IPro | -8.4 | PEC | -1.6 |
| A14C | -14.5 | TPro | -8.3 | VIP | -1.5 |
| A13L | -14.4 | AuR | -8.2 | TLO | -1.2 |
| A13M | -14.3 | AuRM | -8.1 | V5 | -1.1 |
| A47O | -14.0 | A23c | -8.0 | TFO | -1.1 |
| A25 | -13.8 | AuAL | -7.8 | LIP | -0.6 |
| A24a | -13.4 | A3b | -7.7 | PGM | -0.5 |
| A6DR | -13.3 | TPO | -7.6 | V6A | -0.5 |
| A6Vb | -13.2 | Ent | -7.2 | OPt | -0.3 |
| ProM | -13.0 | ReI | -7.1 | A29a-c | -0.3 |
| A6Va | -12.7 | S2I | -7.0 | MIP | -0.1 |
| A8C | -12.7 | A36 | -7.0 | TEO | 0.1 |
| OPAl | -12.6 | S2E | -7.0 | A23V | 0.5 |
| A24b | -12.5 | Area1-2 | -6.8 | ProSt | 0.7 |
| OPro | -12.4 | AuA1 | -6.7 | V4T | 1.1 |
| A24c | -12.2 | A35 | -6.6 | A19M | 1.3 |
| GU | -12.0 | AuCM | -6.4 | V3A | 1.3 |
| A6M | -12.0 | AuCPB | -6.4 | V4 | 2.3 |
| A6DC | -11.7 | AuML | -6.3 | V6 | 2.7 |
| TPPro | -11.2 | PF | -6.1 | A19DI | 3.4 |
| A4c | -11.1 | TE2 | -5.4 | V3 | 4.7 |
| Pir | -11.1 | AuCL | -5.0 | V2 | 5.4 |
| PaIM | -10.8 | PGa/IPa | -4.9 | V1 | 6.6 |
| S2PR | -10.4 | TPt | -4.7 | | |

**Supplementary Table 7 | Rostrocaudal coordinates.** The rostro-caudal coordinate of each of the 116 areas of the marmoset cortex, measured based on the spatial coordinates of their barycenter[9].

**Supplementary Figures**

The macaque data used in Supplementary Figs. 1-3,8-12 are obtained from Markov et. al 2014[10], in Supplementary Figs. 4,15,16 from Chaudhuri et al. 2014[11] and the mouse data used in Supplementary Figs. 9,10 from Gămănuț et al. 2019[12], while for Supplementary Fig. 8 from Horvát et al. 2016[13], where the projection lengths have been reported. We considered connectivity and wiring distances data from these resources for consistency because they have been collected with the same way as the marmoset connectivity data in the current study. The connectivity data (FLN and SLN) in macaque and mouse in these studies have been collected with retrograde tracing, and for the wiring distances the same definition was used (shortest path through the white matter, avoiding the gray matter), and therefore they are suitable for comparison.

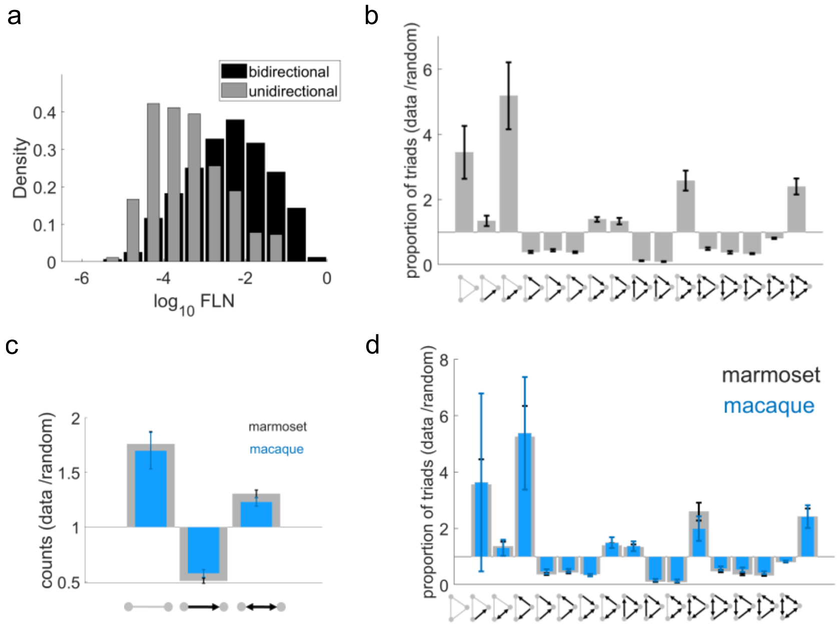


**Supplementary Figure 1 | Two- and three-node motifs in marmoset and macaque cortico-cortical connectivity network. (a)** Distribution of the FLN values of the reciprocal connections of the edge-complete network (black bars; $\mu_{log_{10}FLN_{reciprocal}} = -2.43$) and of the unidirectional connections (gray bars; $\mu_{log_{10}FLN_{unidirectional}} = -3.41$). The two distributions are different (two-sided two-sample Kolmogorov-Smirnov test, $p = 4.39 \times 10^{-75}$ , Hedges' g effect size: $g = 1.02$). **(b)** Average fraction of the proportion of triads in the marmoset network to the proportion of triads in a random network of the same size and same in- and out- degree sequences as in the data, across 100 realization of such random networks (error bars are standard deviations). The three motifs that are mostly overrepresented in the marmoset cortex are those comprised of bidirectional connections and/or absence of connections. **(c)** Average fraction of the two motif counts of the marmoset (gray) and macaque (blue) edge-complete subnetwork to the two motif counts in a randomized version of the edge-complete network keeping the in-and out-degree sequences the same as in the data. The average is across 100 realizations across such random network and the error bars are standard deviations. **(d)** Same as (b) but for the proportion of triads in the marmoset (gray) and macaque (blue) edge complete network.

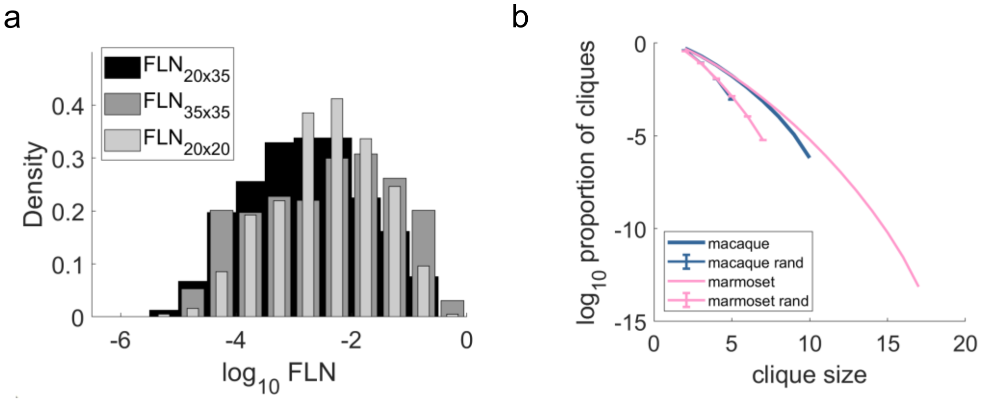


**Supplementary Figure 2 | FLN of the core-periphery subnetworks and cliques in the marmoset and macaque cortico-cortical connectivity network. (a)** Distribution of the FLN values within the core (20 × 20), within the periphery (35 × 35) and between core and periphery (20 × 35). The FLN distributions within the core and periphery come from normal distributions with same mean

($\mu_{log_{10}FLN_{20\times20}} = -2.42$ and $\mu_{log_{10}FLN_{35\times35}} = -2.44$ respectively, two-sided two-sample t-test: $p = 0.8$) but different variance (two-sided two-sample F-test: $p = 5.60 \times 10^{-6}$). Both distributions are different than the distribution of the FLN between core and periphery nodes (two-sided two-sample Kolmogorov-Smirnov test: $p = 4.15 \times 10^{-9}$ , $8.92 \times 10^{-10}$ respectively, Hedges' g effect size: $g = 0.37$, $0.32$ respectively) and tend to have slightly stronger values $\mu_{log_{10}FLN_{20\times35}} = -2.79$). **(b)** Proportion of cliques (how many there are divided by how many they could be based on the size of the edge complete network) of marmoset (pink) and macaque (blue). The plots with the error bars (one standard deviation) are the same counts but averaged across 1000 random networks of same size and keeping the in- and out- degree sequences as in the data.

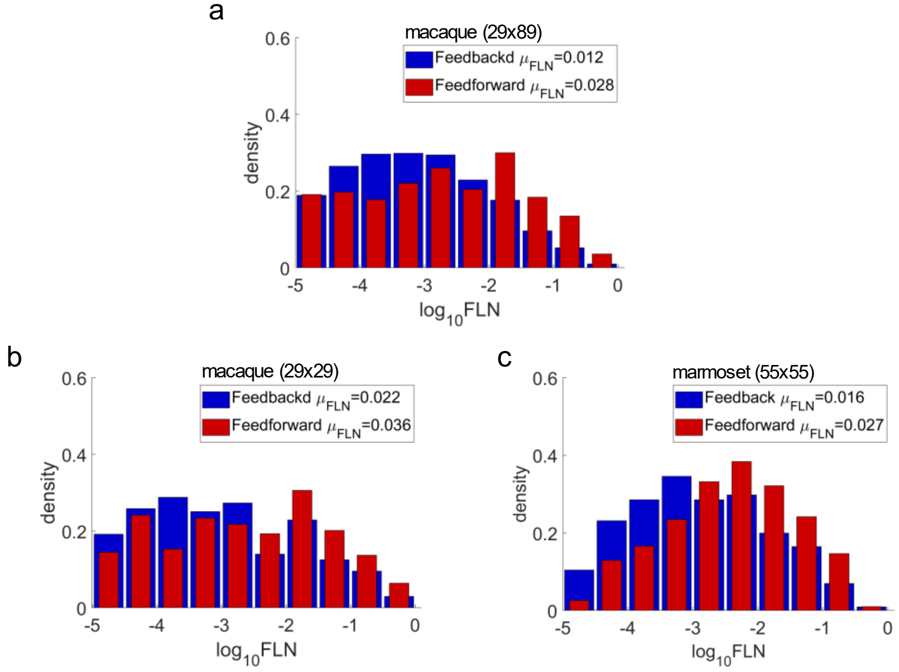


**Supplementary Figure 3 | FLN of feedforward and feedback connections.** Distribution of the FLN values of the feedforward connections (red; SLN > 0. 5) and of the feedback connections (blue; SLN < 0.5), with the first being stronger than the latter of the macaque connectivity 29x89 matrix[11]

(middle), its edge-complete version **(b)**, and of the marmoset edge-complete network **(c)** (two-sided two-sample Kolmogorov-Smirnov test: $p = 0.0034, 8.89 \times 10^{-10}, 3.30 \times 10^{-16}$, and Hedges' g effect size $g = 0.26, 0.30, 0.44$ respectively.

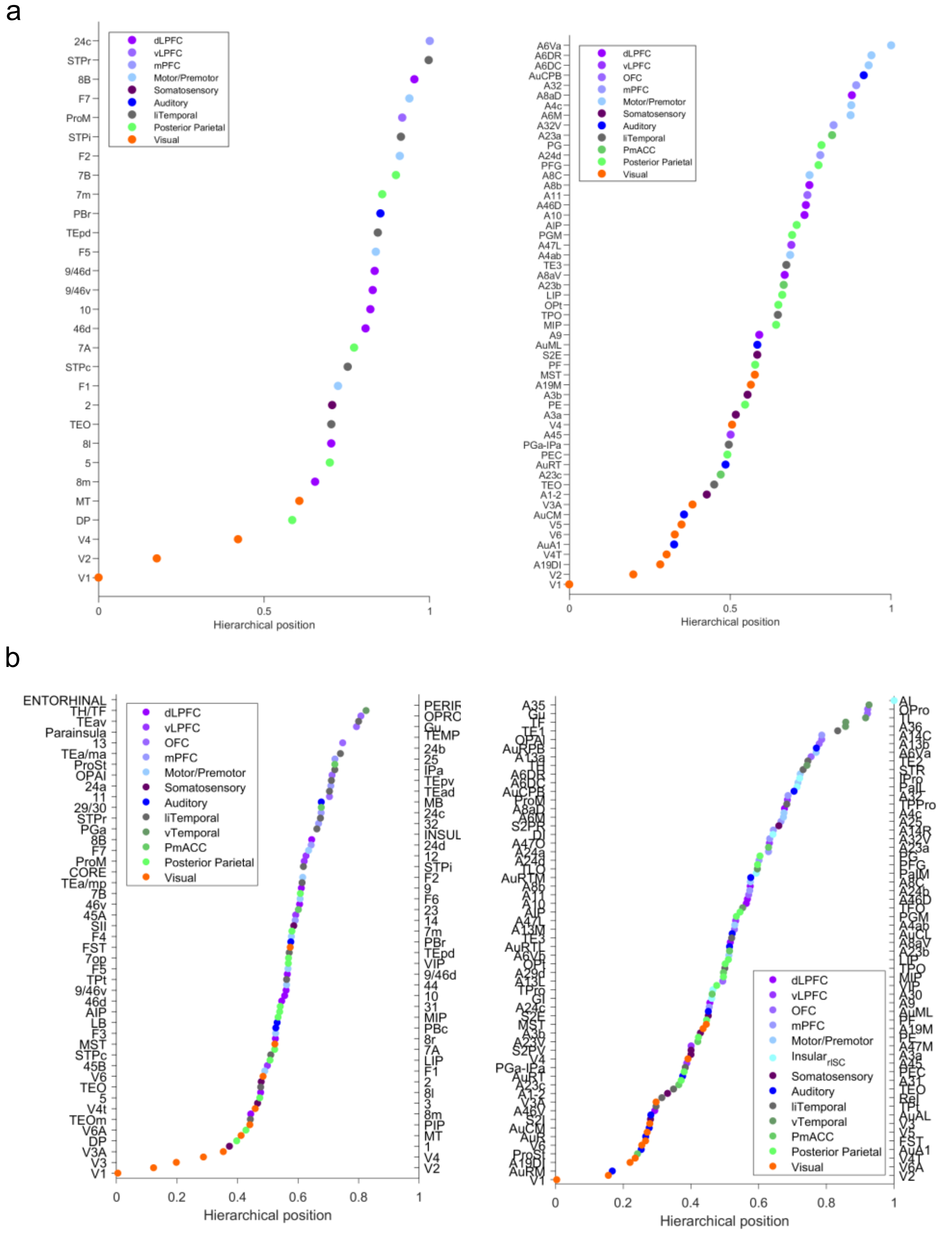


**Supplementary Figure 4 | Hierarchy of marmoset and macaque cortical areas. (a)** Hierarchy of the macaque[11] (left) and marmoset (right) target cortical areas. **(b)** Same as (a) for all the areas.

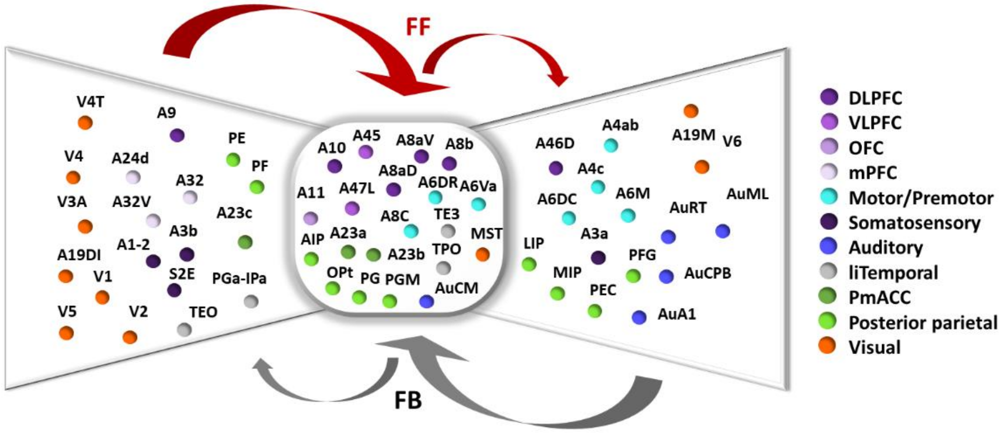


**Supplementary Figure 5 | Bow-tie representation.** Bow-tie representation of the relation of the core structure (Fig. 2e,right) with the periphery, derived from the hierarchical laminar weights of their pathways. FF: feedforward, FB: feedback. Visual, somatosensory and medial prefrontal areas mainly project to the core in an effectively feedforward way while the core projects effectively feedback mainly to auditory, motor and parietal areas. The bow-tie representation s computed by the feedforward and feedback classification of the connections based on the SLN and the core-periphery structure of the network, following the method described in [Markov et al. 2013][14]. In the center of the scheme there is the dense core structure (Fig. 2e,right) while at the wings there are the areas of the periphery (the rest of the target areas) categorized to the left or to the right based on the feedforward/feedback connections into and from the core.

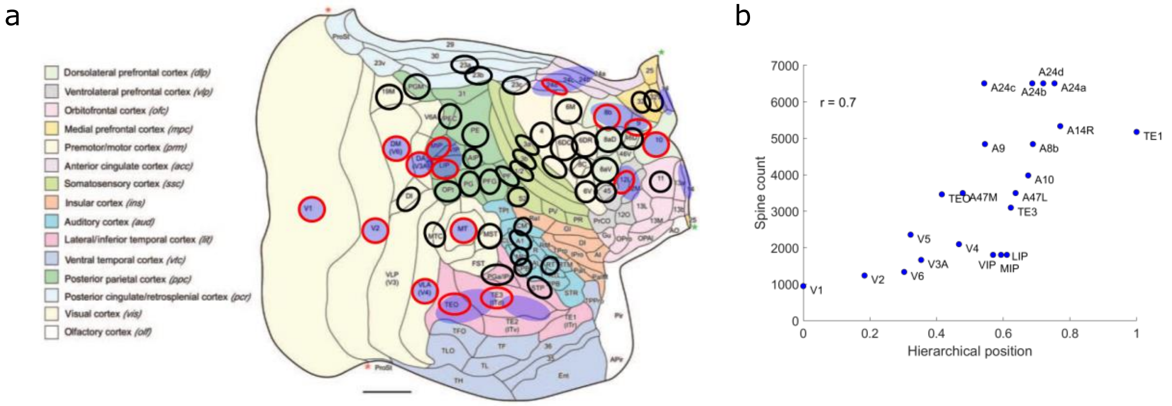


**Supplementary Figure 6 | Spine count as a function of hierarchy. (a)** Target areas (ellipses with black, and red, perimeter) visualized on the marmoset cortical 2D flat map [Reser et al. 2013][15]. Ellipses with blue shaded area correspond to the areas where intracellular injections of fluorescent tracers have been conducted (14 areas in total from which 15 overlap with the target areas denoted as ellipses with red perimeter) and spine count data have been collected from the literature (Supplementary Table 4). **(b)** The spine count of the areas in the Paxinos et al. 2012 parcellation are plotted (Supplementary Table 4) as a function their hierarchical value (Fig. 4a). Here for areas that are merged in the parcellation considered in the source study, the same spine count is assumed.

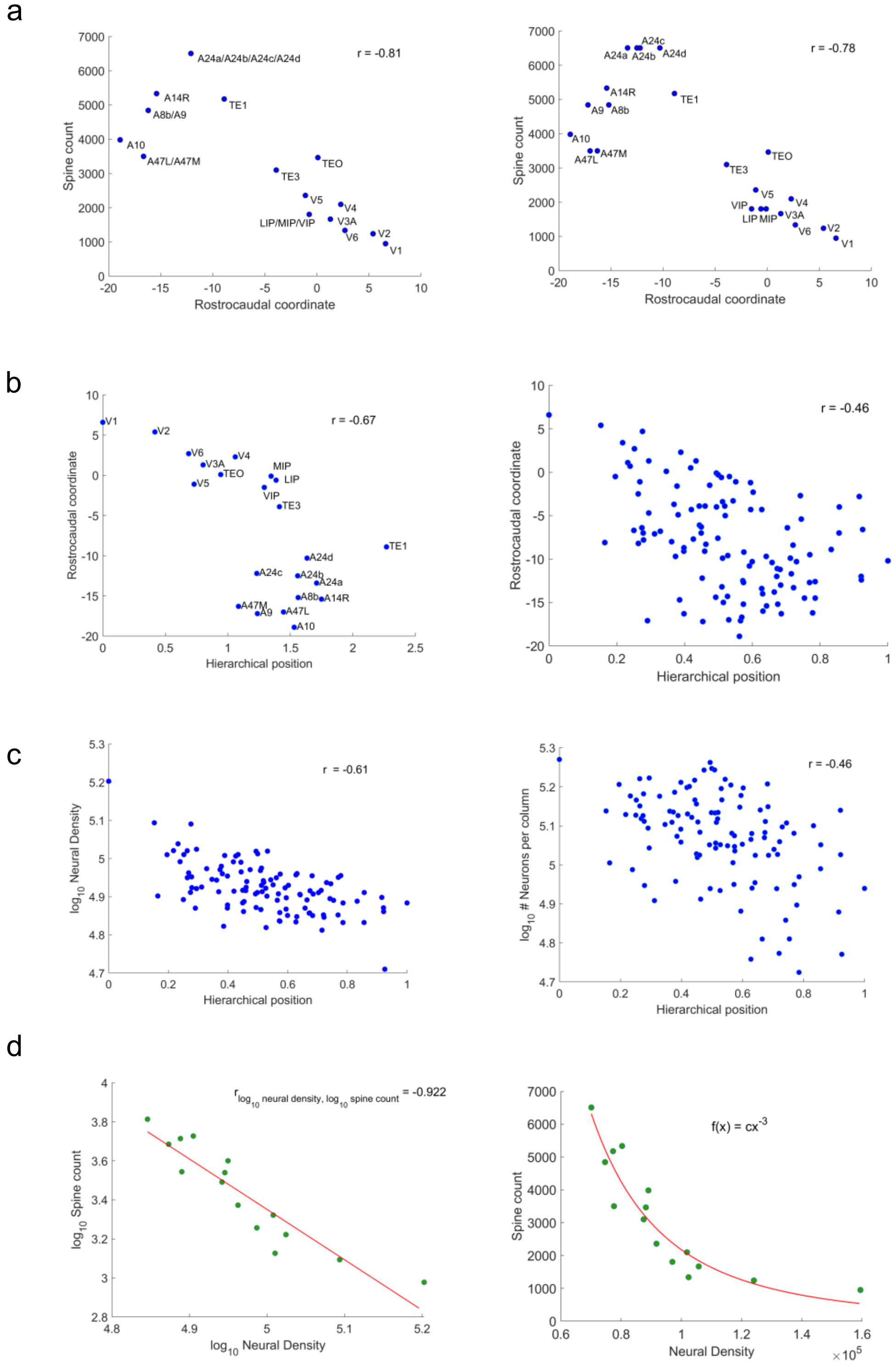
a
Spine count
Rostrocaudal coordinate
r = -0.81
A24a/A24b/A24c/A24d
A14R
A8b/A9
TE1
A10
A47L/A47M
TEO
TE3
V5
V4
LIP/MIP/VIP
V3A
V6
V2
V1
r = -0.78
A24c
A24a
A24b
A24d
A9
A8b
A47M
A47L
VIP
LIP
MIP
b
Rostrocaudal coordinate
Hierarchical position
r = -0.67
r = -0.46
c
$\log_{10}$ Neural Density
r = -0.61
$\log_{10}$ # Neurons per column
r = -0.46
d
$\log_{10}$ Spine count
$\log_{10}$ Neural Density
$r_{\log_{10}\text{ neural density}, \log_{10}\text{ spine count}}$ = -0.922
Spine count
Neural Density
$\times 10^5$
$f(x) = cx^{-3}$

**Supplementary Figure 7 | Microstructural properties as a function of hierarchy. (a)** Left: Spine count of basal dendrite in a layer 3 pyramidal neuron is correlated with the rostrocaudal axis. The rostrocaudal coordinate of the areas that are merged is the average over them (as Fig. 5). Right: Same as left but for the spine count of the areas in the Paxinos et al. 2012 parcellation[5] (as Supplementary Fig. 6b). **(b)** Rostrocaudal coordinates as a function of the hierarchical level of the areas for which we have spine count (as in a,right) and of all areas (right). **(c)** Neural density as a function of the hierarchical level of all areas (left) and number of neurons per column as a function of hierarchy (right). **(d)** Spine count as a function of the common logarithm of the the neural density (left). The red line is linear fit, which gives $log_{10}(spine\ count) = -2.6\ log_{10}(neural\ density) + 16.3$, which is approximated to a cubic power law (right, red line, where the constant c is fitted to the data). The neural density data was obtained from Table 1 in Atapour et al. 2019[9]. The number of neurons per unit column of each marmoset cortical area was taken from Table 3 in [Atapour et al. 2019][9]. The rostrocaudal coordinate of each area is computed based on the stereotaxic coordinates of their barycenter[9] and shown in Supplementary Table 7. *r* is the Pearson correlation.

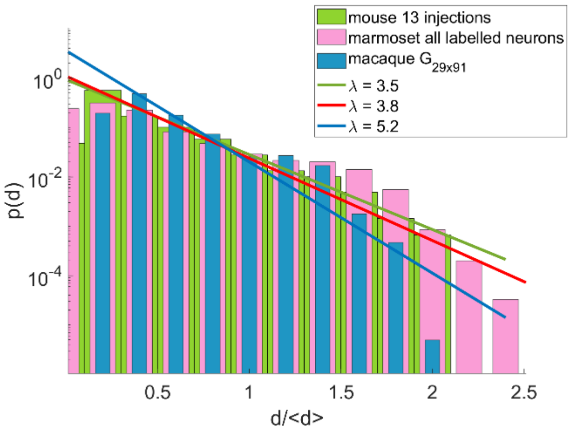


**Supplementary Figure 8 | EDR at the common template.** The exponential distance rule in the common template of mouse (green), marmoset (pink) and macaque (blue). Distance in the x-axis are the wiring distances divided by the average distance as suggested in Horvat et al. 2016[13]. In the marmoset case, all 1,966,028 projection lengths were divided by 12.4375 mm (the marmoset average interareal wiring distance). The bar plots and the corresponding line fits were done as in Fig. 6c. The bin size here is 0.2.

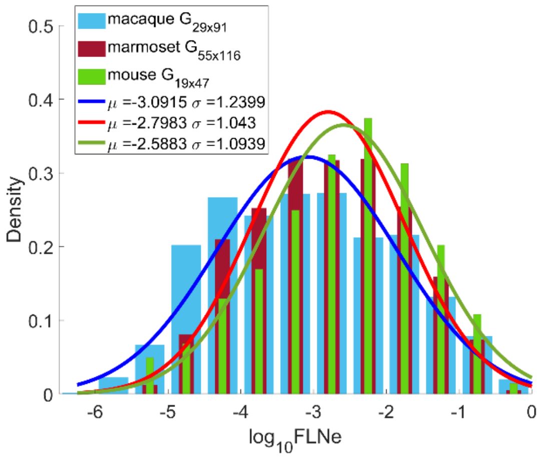


**Supplementary Figure 9 | FLN distribution across species.** The mean of the distribution of the marmoset connectivity weights (red) lies between that of the macaque (blue) and of the mouse (green). Bars are binned data and lines are fits of gaussian distribution with mean and standard deviation from the data. Data from all species come from different distributions (two-sided two-sample Kolmogorov-Smirnov test: $p = 9.93 \times 10^{-10}$, $2.84 \times 10^{-18}$, $p = 5.40 \times 10^{-23}$ and Hedges' g effect size: $g = 0.20$, 0.26, 0.42 for marmoset vs mouse, macaque vs. marmoset and marmoset vs. mouse data respectively. Only the distributions of the marmoset and mouse have similar variances, two-sided two-sample F-test: $p = 0.06$).

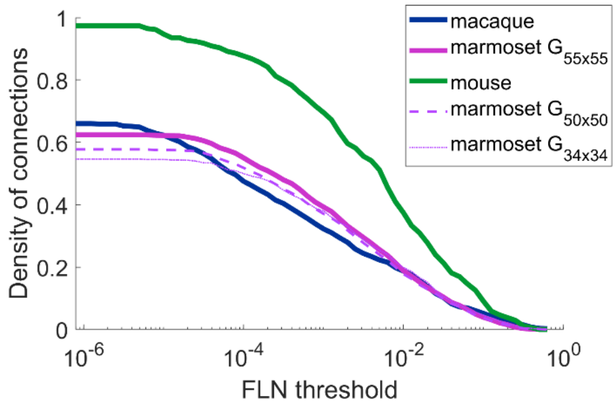


**Supplementary Figure 10 | Density as a function of FLN threshold, across species.** The density of the edge-complete marmoset (purple), macaque (blue) and mouse (green) cortico-corticortical networks as a function of FLN threshold. The data in both primates indicate a same density of the cortical connectome, which is lower than that in the mouse. Different plots for the marmoset correspond to different subsets of the data based on the leakage of the tracer in neighboring areas (see Discussion, Supplementary Fig. 14).

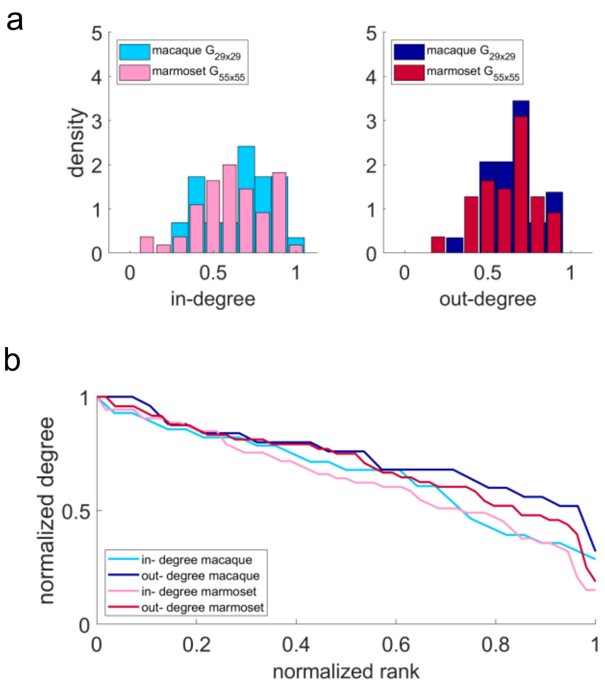

**Supplementary Figure 11 | In- and out- degree of marmoset and macaque cortical areas. (a)** In- and out- degree distribution of the marmoset (pink and red respectively) and macaque (light and dark blue respectively), normalized by the size of the edge-complete network (n = 55 of the marmoset and n = 29 of the macaque). **(b)** Normalized (by the maximum value) in- and out- degree sequences in descending order for macaque and marmoset with color codes as in (a).

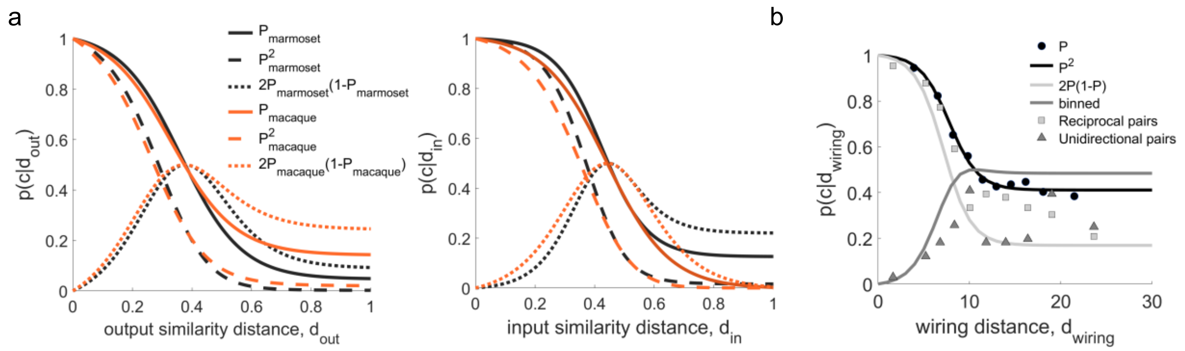


**Supplementary Figure 12 | Probability of connections dependence on functional similarity and wiring distance. (a)** Proportion of connections as a function of similarity distance (left: input similarity distance, right: output similarity distance) for the marmoset (black) and macaque (orange). Here only the plots from fitting on the data is shown (solid lines) and the predicted plots for the reciprocal and non-reciprocal connections (dashed plots) (for details of methods see Song et al. 2014[16]). **(b)** Proportion of connections as a function of wiring distance for the marmoset, as was done for macaque in Song et al. 2014[16].

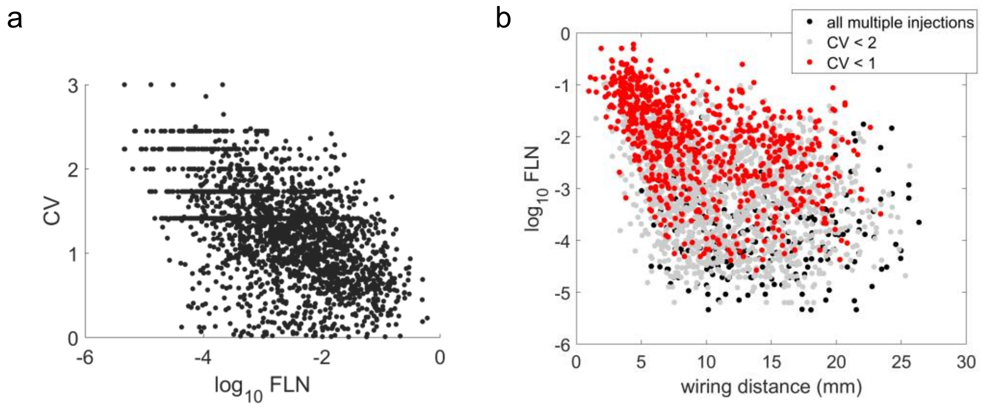


**Supplementary Figure 13 | Variability across injections. (a)** Coefficient of variation (CV) of the FLN as a function of the common logarithm of the average value of the FLN. **(b)** The common logarithm of the FLN as a function of interareal wiring distance for different CV of the FLN thresholds.

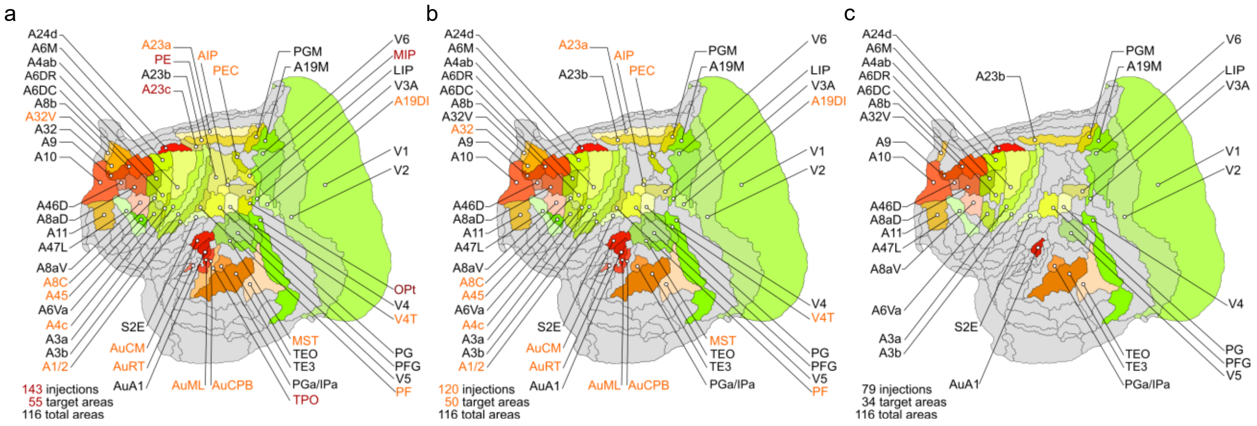


**Supplementary Figure 14 | Samples of data based on the leakage of tracers in adjacent areas.** From the 143 retrograde tracing injections (55 target areas, shown on the 2d flattened map of the marmoset cortex, **a**) in 120 of them the tracer was confinded at least 80% of its overall volume within the injected target area (50 target areas, **b**), and in 79 of them 100% of the tracer's volume was within the target area (34 target areas, **c**) (see supplementary table S3 in Majka et al. 2020[17]).

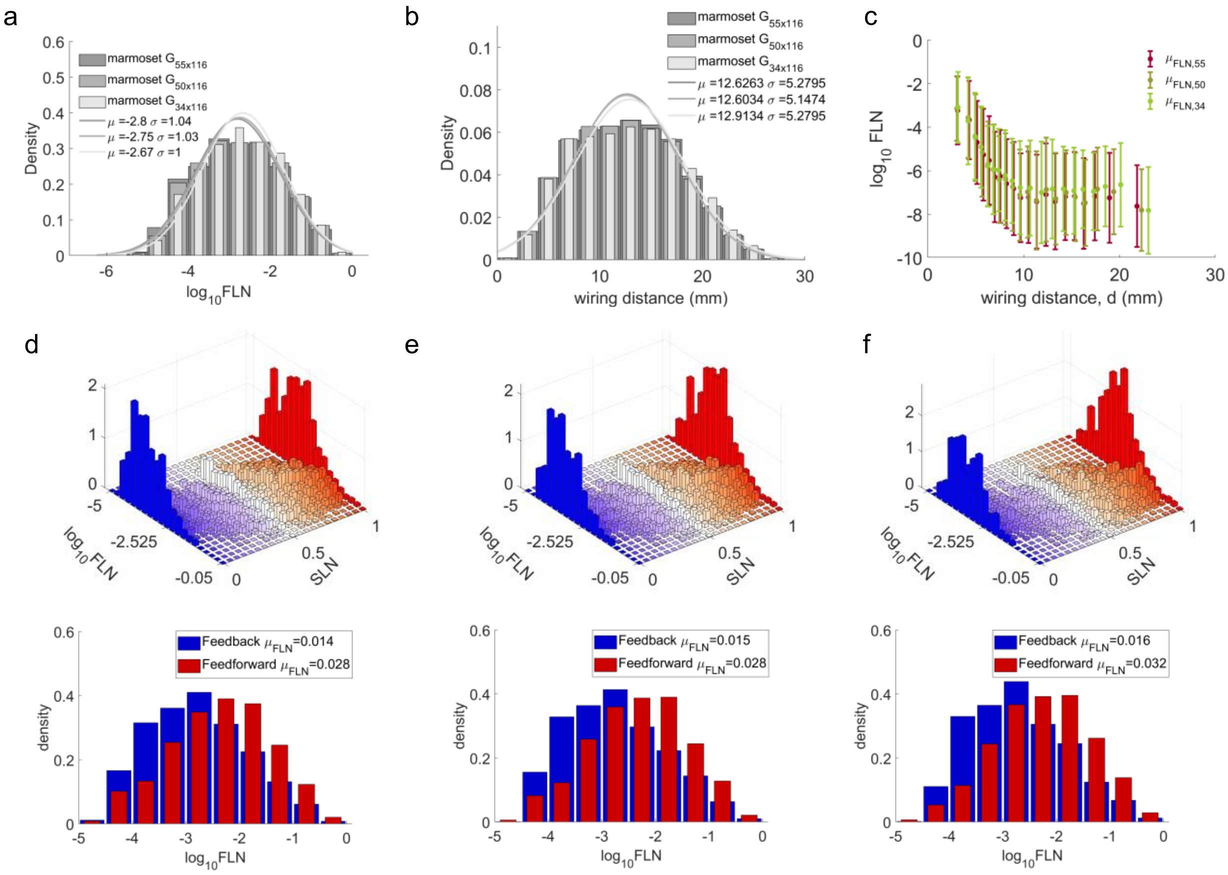


**Supplementary Figure 15 | Analysis of samples of data based on the leakage of tracers in adjacent areas. (a)** Distribution of FLN as in Fig. 1d for the different data subsets based on the leakage of the tracer in neighboring areas (Supplementary Fig. 14). The data for the 55 and 50 target areas come from the same distribution (two-sided two-sample Kolmogorov-Smirnov test： $p = 0.44$), while the data for the 50 and 34 target areas are from two distributions of different means (two-sided two-sample t-test: $p = 0.009$) and same variance (two-sided two-sample F-test: $p = 0.13$) (two-sided two-sample Kolmogorov-Smirnov test： $p = 0.014$, Hedges' g effect size: $g = 0.079$), while that of the 55 and 34 target areas are from different distributions, with small though effect size (two-sided two-sample Kolmogorov-Smirnov test： $p = 3.50 \times 10^{-4}$, Hedges' g effect size: $g = 0.13$). **(b)** Distribution of interareal wiring distances as in Fig. 6a fot the different cases as in (a). The data for the 55, 50, and 34 arget areas come from the same distribution (two-sided two-sample Kolmogorov-Smirnov test： $p = 0.9999$ (for $G_{55\times116}$, $G_{50\times116}$), 0.0821 (for $G_{50\times116}$, $G_{34\times116}$), 0.0515 (for $G_{55\times116}, G_{34\times116}$)). **(c)** The average values (and their standard deviations) of the FLN as a function of wiring distance as shown in Fig. 6b for the different cases as in (a) (cicles and error bars are mean and one standard deviation over a window of 173, 146 and 90 data points respectively, so that the bin size is 20 in all cases). (**d)-(e)**. Up. The 2d histogram of the FLN and SLN for the 55 target areas as in Fig. 3c for the 55, 50 and 34 target areas respectively. Bottom. The distribution of the connectivity weights that correspond to feedforward and feedback projections as in Fig. 3d for the 55, 50 and 34 target areas respectively. The values from all projections (not the average values across injections in the same target areas) are ploted here, but the results don't change if consider the average values and even the corresponding edge-complete matrices. . In all cases the FLN distributions that correspond to feedforward and feedback are different with different mean but same variance (two-sided two-sample Kolmogorov-Smirnov test: $p = 3.87 \times 10^{-65}$, $4.31 \times 10^{-57}$, $2.58 \times 10^{-40}$, Hedges' g effect size $g = 0.46, 0.48, 0.52$, two-sided two-sample t-test: $p = 7.68 \times 10^{-73}$, $4.95 \times 10^{-62}$, $1.93 \times 10^{-43}$, and two-sided two-sample F-test: p = 0.36, 0.63, 0.86 for (d)-(e) respectively).

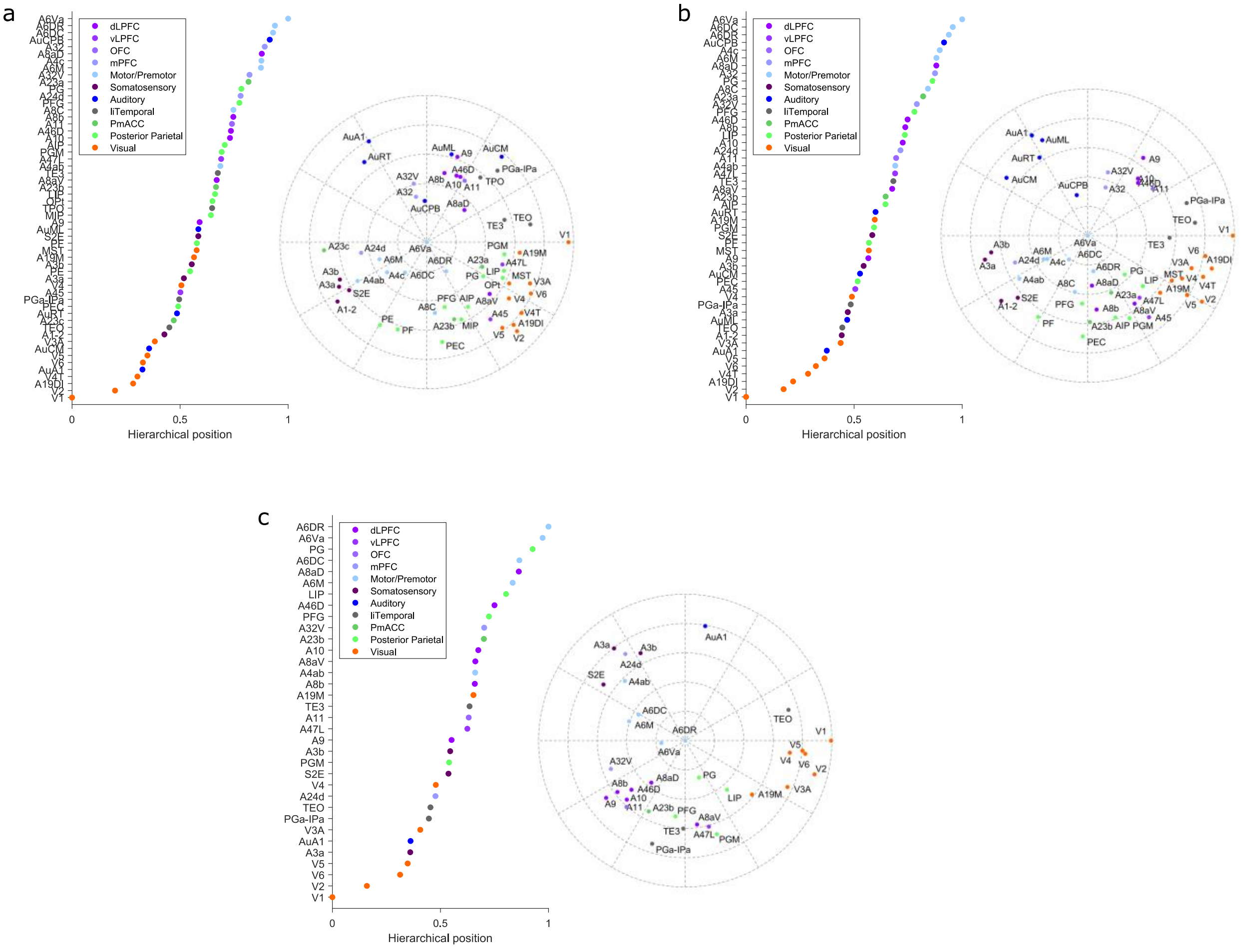


**Supplementary Figure 16 | Hierarchy and two-dimensional representation of the marmoset architecture based on data samples according to the leakage of tracers in adjacent areas.** Hiearchical position and 2d polar plot (insets) as shown in Fig. 4 of the 55 **(a)** 50 **(b)** and 34 **(c)** target areas (Supplementary Fig. 14) computed from their corresponding FLN and SLN data.

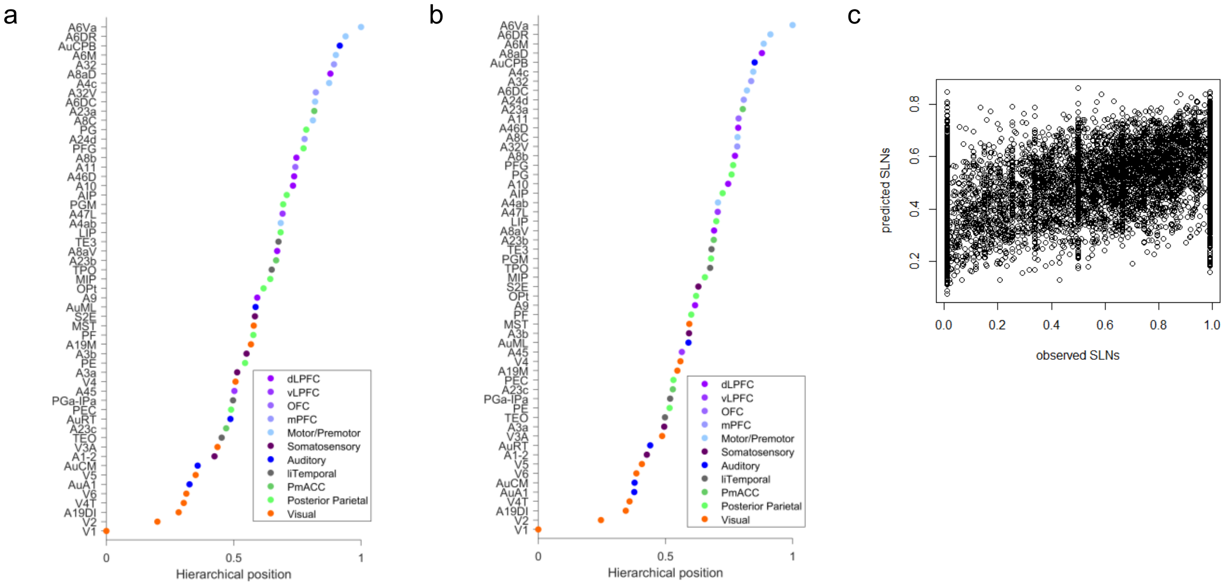


**Supplementary Figure 17 | Comparing hierarchy computed with logistic regression. (a)** Hierarchy of the marmoset cortical areas fitted with beta regression**. (b)** Hierarchy of the marmoset cortical areas fitted with logistic regression. **(c)** Predicted SLN values as a function of the actual SLN values, the Pearson correlation is 0.53.

### References


1. Elston, G. N., Tweedale, R. & Rosa, M. G. P. Cellular heterogeneity in cerebral cortex: a study of the morphology of pyramidal neurones in visual areas of the marmoset monkey. *J. Comp. Neurol.* **415**, 33–51 (1999).
2. Elston, G. N., Benavides-Piccione, R. & DeFelipe, J. The pyramidal cell in cognition: a comparative study in human and monkey. *J. Neuronsci.* **21**, RC163–RC163 (2001).
3. Aoi, H., Oga, T., Sasaki, T., Fujita, I. & Ichinohe, N. Postnatal development of layer III pyramidal cells in the primary visual, inferior temporal, and prefrontal cortices of the marmoset. *Front. Neural Circuits* **7**, 1–10 (2013).

4. Sasaki, E. Prospects for genetically modified non-human primate models, including the common marmoset. *Neurosci. Res.* **93**, 110–115 (2015).
5. Paxinos, G., Watson, C., Petrides, M., Rosa, M. & Tokuno, H. *The marmoset brain in stereotaxic coordinates*. (Academic Press, 2012).
6. Hofman, M. A. Size and shape of the cerebral cortex in mammals. II. The cortical volume. *Brain Behav. Evol.* **32**, 17–26 (1988).
7. Zhang, K. & Sejnowski, T. J. A universal scaling law between gray matter and white matter of cerebral cortex. *PNAS* **97**, 5621–5626 (2000).
8. Navarrete, A. F. *et al.* Primate Brain Anatomy: New Volumetric MRI Measurements for Neuroanatomical Studies. *Brain Behav. Evol.* **91**, 109–117 (2018).
9. Atapour, N. *et al.* Neuronal distribution across the cerebral cortex of the marmoset monkey (callithrix jacchus). *Cereb. Cortex* **29**, 3836–3863 (2019).
10. Markov, N. T. *et al.* Weight consistency specifies regularities of macaque cortical networks. *Cereb. Cortex* **21**, 1254–1272 (2011).
11. Chaudhuri, R., Knoblauch, K., Gariel, M.-A., Kennedy, H. & Wang, X.-J. A large-scale circuit mechanism for hierarchical dynamical processing in the primate cortex. *Neuron* **88**, 419–431 (2015).
12. Gămănuţ, R. *et al.* The mouse cortical connectome, characterized by an ultra-dense cortical graph, maintains specificity by distinct connectivity profiles. *Neuron* **97**, 698-715.e10 (2018).
13. Horvát, S. *et al.* Spatial embedding and wiring cost constrain the functional layout of the cortical network of rodents and primates. *PLoS Biol.* **14**, e1002512 (2016).
14. Markov, N. T. *et al.* Cortical high-density counterstream architectures. *Science* **342**, 1238406–1238406 (2013).

15. Reser, D. H. *et al.* Contrasting Patterns of Cortical Input to Architectural Subdivisions of the Area 8 Complex: A Retrograde Tracing Study in Marmoset Monkeys. *Cereb. Cortex* **23**, 1901–1922 (2013).

16. Song, H. F., Kennedy, H. & Wang, X.-J. Spatial embedding of structural similarity in the cerebral cortex. *PNAS* **111**, 16580–16585 (2014).

17. Majka, P. *et al.* Open access resource for cellular-resolution analyses of corticocortical connectivity in the marmoset monkey. *Nature Communications* **11**, 1–14 (2020).
